# Biophysical mechanisms in the mammalian respiratory oscillator re-examined with a new data-driven computational model

**DOI:** 10.1101/415190

**Authors:** Ryan S. Phillips, Tibin T. John, Hidehiko Koizumi, Yaroslav I. Molkov, Jeffrey C. Smith

## Abstract

An autorhythmic population of excitatory neurons in the brainstem pre-Bötzinger complex is a critical component of the mammalian respiratory oscillator. Two intrinsic neuronal biophysical mechanisms—a persistent sodium current (*I*_*NaP*_) and a calcium-activated non-selective cationic current (*I*_*CAN*_)—were proposed to individually or in combination generate cellular-and circuit-level oscillations, but their roles are debated without resolution. We re-examined these roles with a new computational model of an excitatory population with randomly distributed *I*_*NaP*_ and *I*_*CAN*_ conductances and synaptic connections. This model robustly reproduces experimental data showing contrary to previous hypotheses, rhythm generation is independent of *I*_*CAN*_ activation, which instead determines population activity amplitude. The novel insight is that this occurs when *I*_*CAN*_ is primarily activated by neuronal calcium fluxes driven by synaptic mechanisms. Rhythm depends critically on *I*_*NaP*_ in a subpopulation forming the rhythmogenic kernel. The model explains how the rhythm and amplitude of respiratory oscillations involve distinct biophysical mechanisms.

## Introduction

Defining cellular and circuit mechanisms generating the vital rhythm of breathing in mammals remains a fundamental unsolved problem of wide-spread interest in neurophysiology (Richter & Smith, 2014; Del Negro et al., 2018), with potentially far-reaching implications for understanding mechanisms of oscillatory circuit activity and rhythmic motor pattern generation in neural systems (Marder & Calabrese, 1996; Buzsaki, 2006; Grillner, 2006; Kiehn, 2006). The brainstem pre-Bötzinger complex (pre-BötC) region (Smith et al., 1991) located in the ventrolateral medulla oblongata is established to contain circuits essential for respiratory rhythm generation (Smith et al., 2013; Del Negro et al., 2018), but the operational cellular biophysical and circuit synaptic mechanisms are continuously debated. Pre-BötC excitatory neurons and circuits have autorhythmic properties and drive motor circuits that can be isolated and remain rhythmically active in living rodent brainstem slices in vitro. Numerous experimental and theoretical analyses have focused on the rhythmogenic mechanisms operating in these in vitro conditions to provide insight into biophysical and circuit processes involved, with potential relevance for rhythm generation during breathing in vivo (Feldman & Del Negro, 2006; Richter & Smith, 2014). The ongoing rhythmic activity in vitro has been suggested to arise from either a subset(s) of intrinsically bursting neurons which, through excitatory synaptic interactions, recruit and synchronize neurons within the network (pacemaker-network model) (Rekling & Feldman, 1998; Toporikova & Butera, 2011; Ramirez et al., 2004), or as an emergent network property through recurrent excitation (Jasinski et al., 2013) and/or synaptic depression (group pacemaker model) (Rubin et al., 2009).

From these previous analyses, involvement of two possible cellular-level biophysical mechanisms have been proposed. One based on a slowly inactivating persistent sodium current (*I*_*NaP*_) (Butera et al., 1999a), and the other on a calcium-activated non-selective cation current (*I*_*CAN*_) coupled to intracellular calcium ([*Ca*]_*i*_) dynamics (for reviews see (Rybak et al., 2014; Del Negro et al., 2010), or a combination of both mechanisms (Jasinski et al., 2013). Despite the extensive experimental and theoretical investigations of these sodium-and calcium-based mechanisms, the actual roles of *I*_*NaP*_, *I*_*CAN*_ and the critical source(s) of [*Ca*]_*i*_ transients in the pre-BötC are still unresolved. Furthermore, in pre-BötC circuits the process of rhythm generation must be associated with an amplitude of circuit activity sufficient to drive downstream circuits to produce adequate inspiratory motor output. Biophysical mechanisms involved in generating the amplitude of pre-BötC circuit activity have also not been established.

*I*_*NaP*_ is proposed to mediate an essential oscillatory burst-generating mechanism since pharmacologically inhibiting *I*_*NaP*_ abolishes intrinsic neuronal rhythmic bursting as well as pre-BötC circuit inspiratory activity and rhythmic inspiratory motor output in vitro. Theoretical models of cellular and circuit activity based on *I*_*NaP*_ - dependent bursting mechanisms closely reproduce experimental observations such as voltage-dependent frequency control, spike-frequency adaptation during bursts, and pattern formation of inspiratory motor output (Butera et al., 1999b; Pierrefiche et al., 2004; Smith et al., 2007). This indicates the plausibility of *I*_*NaP*_-dependent rhythm generation.

In the pre-BötC, *I*_*CAN*_ was originally postulated to underlie intrinsic pacemaker-like oscillatory bursting at the cellular level and contribute to circuit-level rhythm generation, since intrinsic bursting in a subset of neurons in vitro was found to be terminated by the *I*_*CAN*_ inhibitor flufenamic acid (FFA) (Pena et al., 2004). Furthermore, inhibition of *I*_*CAN*_ in the pre-BötC reduces the amplitude of the rhythmic depolarization (inspiratory drive potential) driving neuronal bursting and can eliminate inspiratory motor activity in vitro (Pace et al., 2007). *I*_*CAN*_ became the centerpiece of the group pacemaker model for rhythm generation, in which this conductance was proposed to be activated by inositol trisphosphate (IP3) receptor/ER-mediated intracellular calcium fluxes initiated via glutamatergic metabotropic receptor-mediated signaling in the pre-BötC excitatory circuits. The molecular correlate of *I*_*CAN*_ was postulated to be the transient receptor potential channel M4 (TRPM4) (Mironov 2008; Pace et al. 2007)– one of the two known Ca^2+^-activated TRP channels (Guinamard et al., 2010; Ullrich et al., 2005). TRPM4 has now been identified by immunolabeling and RNA expression profiling in pre-BötC inspiratory neurons in vitro (Koizumi et al., 2018).

Investigations into the sources of intracellular Ca^2+^ activating *I*_*CAN*_/TRPM4 suggested that (1) somatic calcium transients from voltage-gated sources do not contribute to the inspiratory drive potential (Morgado-Valle et al., 2008), (2) IP3/ER-mediated intracellular Ca^2+^ release does not contribute to inspiratory rhythm generation in vitro), and (3) in the dendrites calcium transients may be triggered by excitatory synaptic inputs and travel in a wave propagated to the soma (Mironov 2008). Theoretical studies have demonstrated the plausibility of [*Ca*]_*i*_-*I*_*CAN*_ dependent bursting (Rubin et al., 2009; Toporikova & Butera, 2011), however these models omit *I*_*NaP*_ and/or depend on additional unproven mechanisms to generate intracellular calcium oscillations to provide burst termination, such as IP3 dependent calcium-induced calcium release (Toporikova & Butera, 2011), partial depolarization block of action potentials (Rubin, et al., 2009), and the Na^+^/K^+^ pump (Jasinski et al., 2013). Surprisingly, pharmacological inhibition of *I*_*CAN*_/TRPM4 has recently been shown to produce large reductions in the amplitude of pre-BötC inspiratory neuron population activity without significant perturbations of inspiratory rhythm (Koizumi et al., 2018). These new observations re-define the role of *I*_*CAN*_, and require theoretical re-examination of pre-BötC neuronal conductance mechanisms and network dynamics, particularly how rhythm generation mechanisms can be independent of *I*_*CAN*_-dependent mechanisms generating the amplitude of network activity.

In this theoretical study, we examine the role of *I*_*CAN*_ in the pre-BötC by considering two plausible mechanisms of intracellular calcium fluxes: (1) from voltage-gated, and (2) from synaptically activated sources. We deduce that *I*_*CAN*_ is primarily activated by calcium transients that are coupled to rhythmic excitatory synaptic inputs originating from *I*_*NaP*_ dependent bursting inspiratory neurons. Additionally, we show that *I*_*CAN*_ contributes to the inspiratory drive potential by mirroring the excitatory synaptic current. Our new model explains the recent experimental observations obtained from in vitro neonatal rodent slices isolating the pre-BötC, showing large reductions in circuit activity amplitude by inhibiting *I*_*CAN*_/TRPM4 without perturbations of inspiratory rhythm generation in pre-BötC excitatory circuits in vitro. The model supports the novel concept that *I*_*CAN*_ activation in a subpopulation of pre-BötC excitatory neurons are critically involved in amplifying synaptic drive from a subset of neurons whose rhythmic bursting is critically dependent on *I*_*NaP*_ and forms the kernel for rhythm generation in vitro. The model shows how the functions of generating the rhythm and amplitude of inspiratory oscillations in pre-BötC excitatory circuits are determined by distinct biophysical mechanisms.

## Results

### 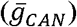 Variation has Opposite Effects on Amplitude and Frequency of Network Bursting in the *Ca*_*V*_ and *Ca*_*syn*_ Models

Recent experimental work (Koizumi et al., 2018) has demonstrated that pharmacological inhibition of *I*_*CAN*_/TRPM4 in the pre-BötC in slices from in vitro neonatal mouse/rat slice preparations, strongly reduces the amplitude of (or completely eliminates) the inspiratory hypoglossal (XII) motor output, as well as the amplitude of pre-BötC excitatory circuit activity that is highly correlated with the decline of XII activity, while having little effect on inspiratory burst frequency. Here, we systematically examine the relationship between *I*_*CAN*_ conductance (*g*_*CAN*_) on amplitude and frequency of circuit activity for voltage-gated (*Ca*_*V*_) and synaptically activated sources (*Ca*_*Syn*_) of intracellular calcium. We found that that reduction of 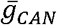 drives opposing effects on circuit activity amplitude and frequency that are dependent on the source of intracellular calcium transients (Fig. 1). In the *Ca*_*V*_ network, where calcium influx is generated exclusively from voltage-gated calcium channels, increasing 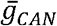 has no effect on amplitude but increases the frequency of network oscillations (Fig. 1A, C, D). Conversely, in the *Ca*_*Syn*_ network where calcium influx is generated exclusively by excitatory synaptic input, increasing 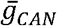 strongly increases the amplitude and slightly decreases the oscillation frequency (Fig. 1B, C, D).

**Figure 1.**
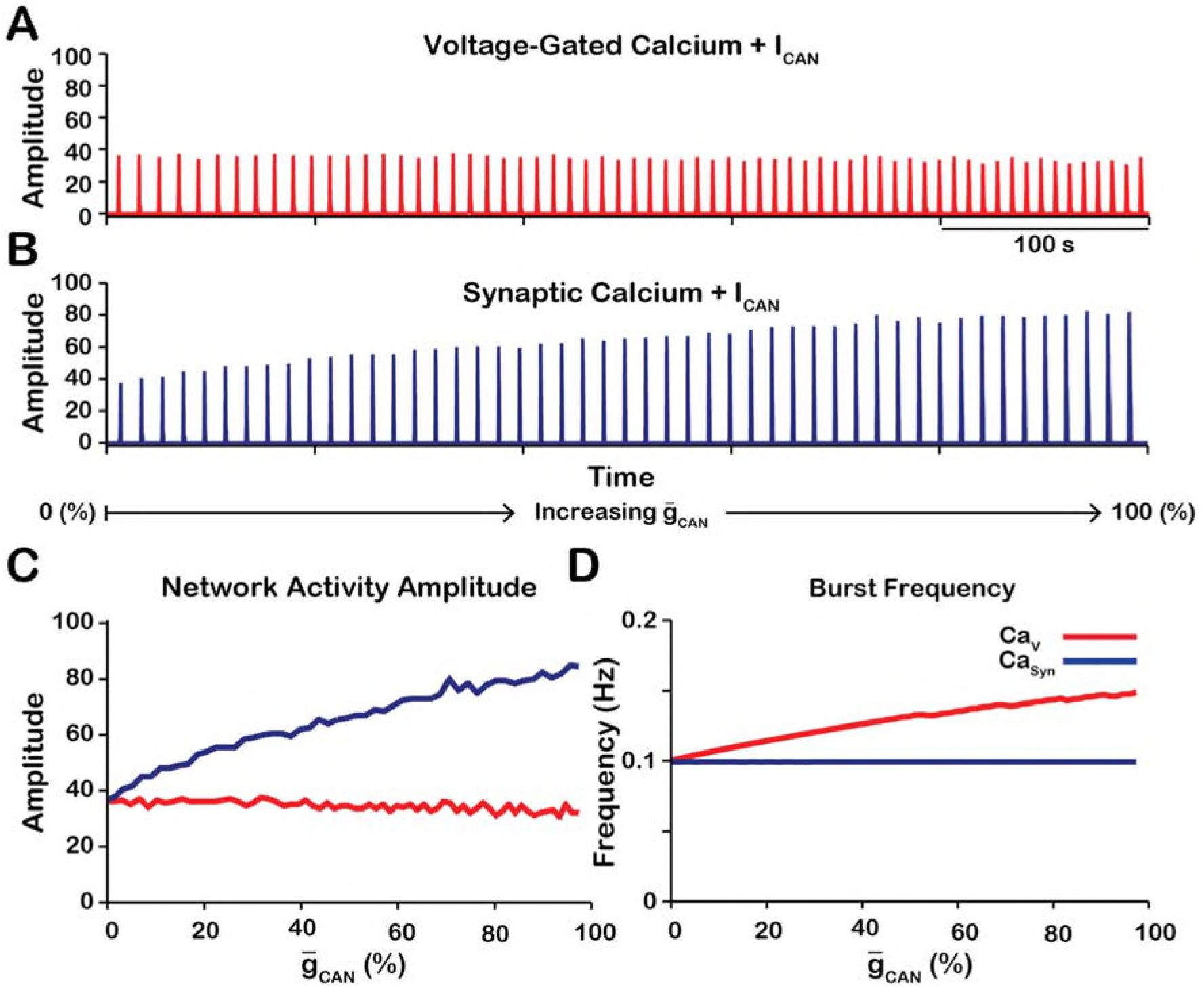
Manipulations of 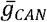 in the *Ca*_*V*_ and *Ca*_*Syn*_ networks produce opposite effects on network activity amplitude (spikes/s) and frequency. **(A & B)** Histograms of neuronal firing and voltage traces for pacemaker and follower neurons in the *Ca*_*V*_, and *Ca*_*Syn*_ networks with linearly increasing 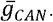 **(C)** Plot of 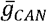 vs network activity amplitude for the *Ca*_*V*_ and *Ca*_*Syn*_networks in **A** and **B**. **(D)** Plot of 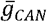 vs network frequency for the *Ca*_*V*_ and *Ca*_*Syn*_ networks in A and B. *Ca*_*V*_ network parameters: 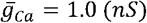, *P*_*Ca*_= 0.0, *P*_*Syn*_= 0.05 and *W*_*max*_*= 0.2 (ns)*. *Ca*_*Syn*_ network parameters: 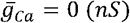, *P*_*Ca*_= 0.01, *P*_*Syn*_= 0.05 and *W*_*max*_= 0.2 (*ns*).

### Effects of Subthreshold Activation of *I*_*CAN*_ on Network Frequency

In *I*_*NaP*_-dependent bursting neurons in the pre-BötC, bursting frequency depends on their excitability (i.e., baseline membrane potential) which can be controlled in different ways, e.g. by directly injecting a depolarizing current (Smith et al., 1991) or varying the conductance and/or reversal potentials of some ionic channels (Butera et al., 1999a). Due to their relatively short duty cycle, the bursting frequency in these neurons is largely determined by the interburst interval, defined as the time between the end of one burst and the start of the next. During the burst, *I*_*NaP*_ slowly inactivates resulting in burst termination and abrupt hyperpolarization of the membrane. The interburst interval is then determined by the amount of time required for *I*_*NaP*_ to recover from inactivation and return the membrane potential back to the threshold for burst initiation. This process is governed by the kinetics of *I*_*NaP*_ inactivation gating variable *h*_*NaP*_. Higher neuronal excitability reduces the value of *h*_*NaP*_ required to initiate bursting. Consequently, the time required to reach this value is decreased, which results in a shorter interburst interval and increased frequency.

To understand how changing 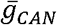 affects network bursting frequency we quantified the values of *h*_*NaP*_ averaged over all pacemaker neurons immediately preceding each network burst and, also, the average *I*_*CAN*_ values between the bursts in the *Ca*_*V*_ and *Ca*_*syn*_ networks (Fig. 2). In the *Ca*_*V*_ network, *I*_*Ca*_ as modeled remains residually activated between the bursts thus creating the background calcium concentration which partially activates *I*_*CAN*_. Therefore, between the bursts *I*_*CAN*_ functions as a depolarizing leak current. Consistently, we found that in the *Ca*_*V*_ network increasing 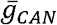 increases *I*_*CAN*_ (Fig. 2A) progressively depolarizing the network, which reduces the *h*_*NaP*_ threshold for burst initiation (Fig. 2B) and, thus, increases network frequency (Fig. 1D).

**Figure 2.**
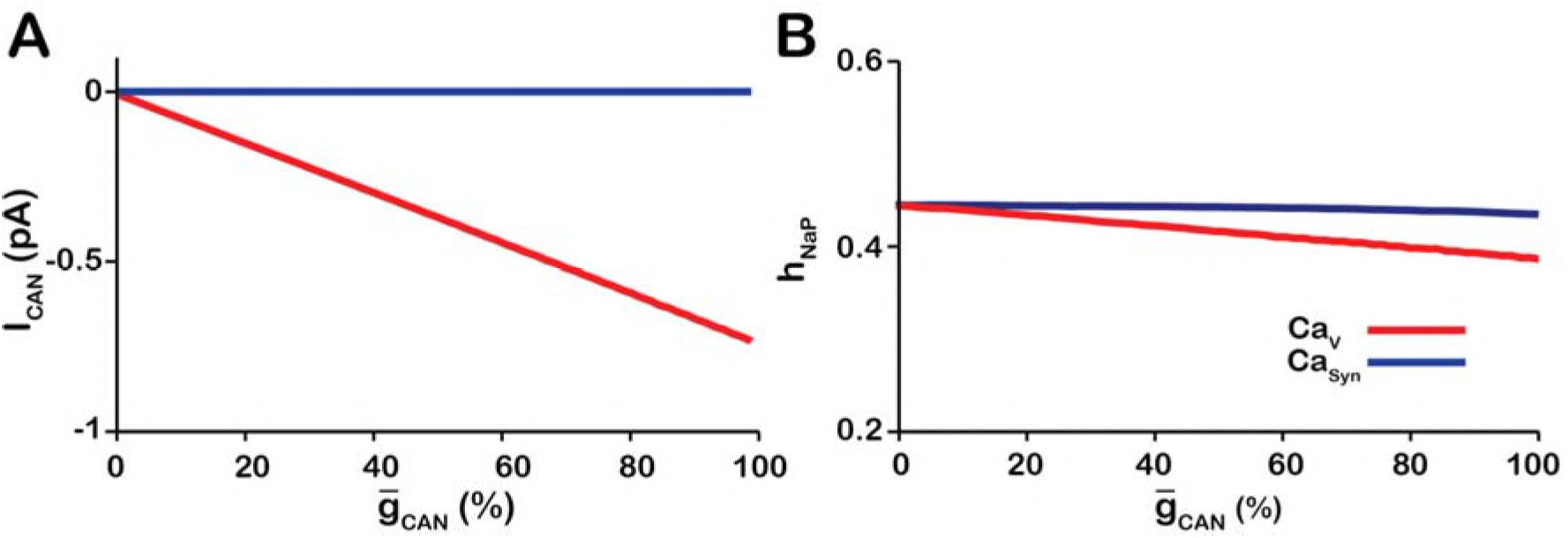
Calcium source and 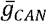-dependent effects on cellular properties regulating network frequency for the simulations presented in Fig. 1. **(A)** Average magnitude of *I*_*CAN*_ in pacemaker neurons during the interburst interval for the *Ca*_*V*_ (red) and *Ca*_*Syn*_ (blue) networks. **(B)** Average inactivation of the burst generating current *I*_*NaP*_ in pacemaker neurons immediately preceding each network burst as a function of 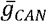 for the voltage-gated and synaptic calcium networks. *Ca*_*V*_ Network Parameters: 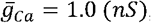, *P*_*Ca*_*= 0.0*, *P*_*Syn*_*= 0.05* and *W*_*max*_*= 0.2 (ns)*. *Ca*_*Syn*_ network parameters: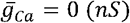, *P*_*Ca*_*= 0.01*, *P*_*Syn*_*= 0.05* and *W*_*max*_ = 0.2 (*ns*).

In the *Ca*_*syn*_ model the intracellular calcium depletes entirely during the interburst interval. Consequently, increasing 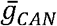 has no effect on *I*_*CAN*_ (Fig. 2A) and frequency is essentially unaffected (Fig. 1D).

### Changes in Network Activity Amplitude Are Driven by Recruitment of Neurons

Network activity amplitude is defined as the total number of spikes produced by the network per a time bin. Consequently, changes in amplitude can only occur by increasing the number of neurons participating in bursts (recruitment) and/or increasing the firing rate of the recruited neurons. To analyze changes in amplitude, we quantified the number of recruited neurons (Fig. 3A) and the average spike frequency in recruited neurons (Fig. 3B) as a function of 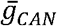for both network models. In the *Ca*_*V*_ network, increasing 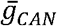 increases the number of recruited neurons (Fig. 3A), but decreases the average spiking frequency in recruited neurons (Fig. 3B) which, together result in no change in amplitude (Fig. 1C). In the *Ca*_*Syn*_ network, increasing 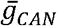 strongly increases the number of recruited neurons (Fig. 3A) and increases the spike frequency of recruited neurons (Fig. 3B) resulting in a large increase in network activity amplitude (Fig. 3C).

**Figure 3.**
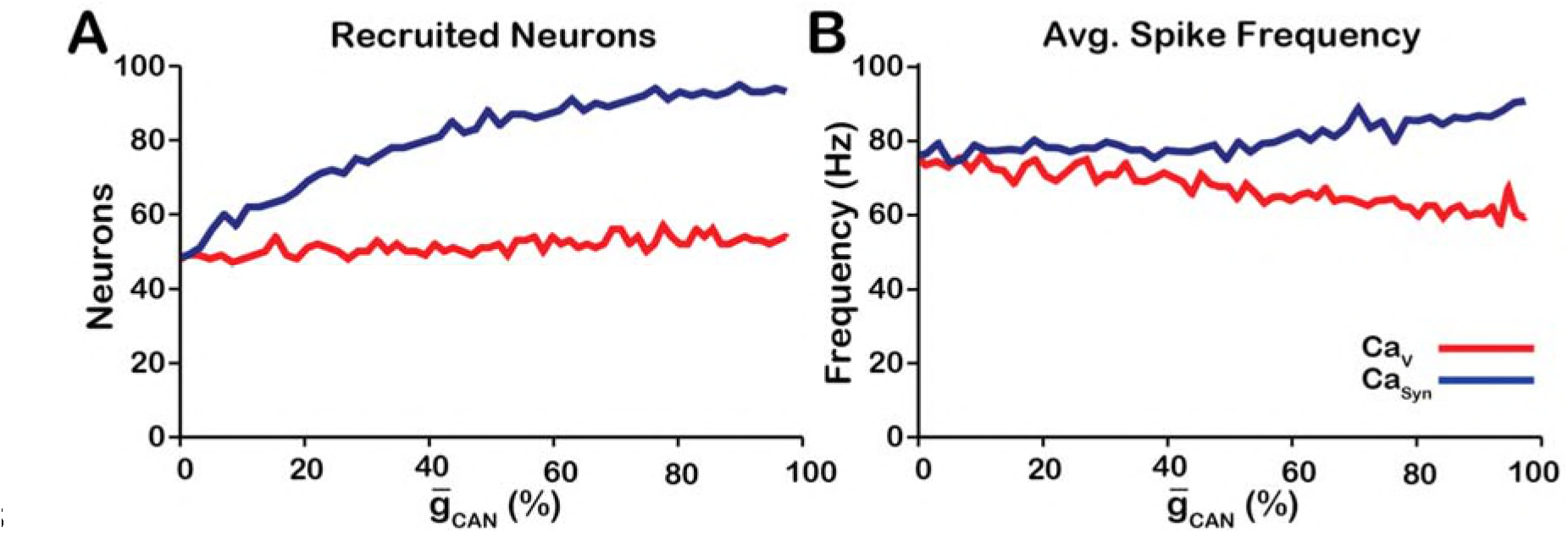
Calcium source and 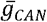-dependent effects on cellular properties regulating network activity amplitude for the simulations presented in Fig. 1. **(A)** Number of recruited neurons in the modeled population of 100 neurons as a function of 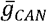for voltage-gated and synaptic calcium sources. The number of recruited neurons is defined as the peak number of spiking neurons per bin during a network burst. **(B)** Average spiking frequency of recruited neurons as a function of 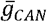 for the voltage-gated and synaptic calcium mechanism. Average spiking frequency is defined the number of spikes per bin divided by the number of recruited neurons. The parameters used in these simulations are: *Ca*_*V*_: 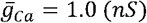, *P*_*Ca*_*= 0.0*, *P*_*Syn*_*= 0.05* and *W = 0.2 (ns)*. *Ca*_*Syn*_: 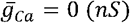*(ns)*, *P*_*Ca*_*= 0.01*, *P*_*Syn*_*= 0.05* and *W= 0.2 (ns)*.

### Manipulating 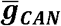 in the *Ca*_*syn*_ Model is Qualitatively Equivalent to Changing the Strength of Synaptic Interactions

Since changes in 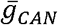 in the *Ca*_*Syn*_ model primarily affect network activity amplitude through recruitment of follower neurons, and the network activity amplitude strongly depends on the strength of synaptic interactions, we next examined the relationship between 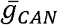, synaptic strength and network activity amplitude and frequency (Fig. 4). Synaptic strength is defined as the number of neurons multiplied by the connection probability multiplied by the average weight of synaptic connections 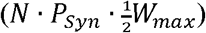. We found that the effects of varying 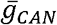 or the synaptic strength on network activity amplitude and frequency are qualitatively equivalent in the *Ca*_*Syn*_ network which is indicated by symmetry of the heat plots (across the X=Y line) in Fig. 4A, B. We further investigated and compared the effect of reducing 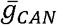 or the synaptic strength on network activity amplitude and frequency as well as the effects on the recruitment of follower neurons (Fig. 4C-F). To make this comparison, picked a starting point in the 2D parameter space between 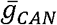 and synaptic strength where the network is bursting. Then in separate simulations we linearly reduced either 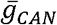 or the synaptic strength to zero. We show that reducing either 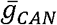 or the synaptic strength have very similar effects on network activity amplitude and frequency (Fig. 4C, D). Furthermore, the effect on follower neurons in both cases is nearly identical (Fig. 4E, F). Reducing either 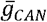 or the synaptic strength decreases the excitatory input to follower neurons during network oscillations which is a major component of the inspiratory drive potential. Therefore, in the *Ca*_*Syn*_ network, manipulations of 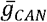 will affect the strength of the inspiratory drive potential in follower neurons in a way that is equivalent to changing the synaptic strength of the network. In contrast, manipulations of 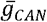 in the *Ca*_*V*_ network will only slightly affect the inspiratory drive potential in follower neurons due to changes in the average firing rate of active neurons (see Fig. 3B).

**Figure 4.**
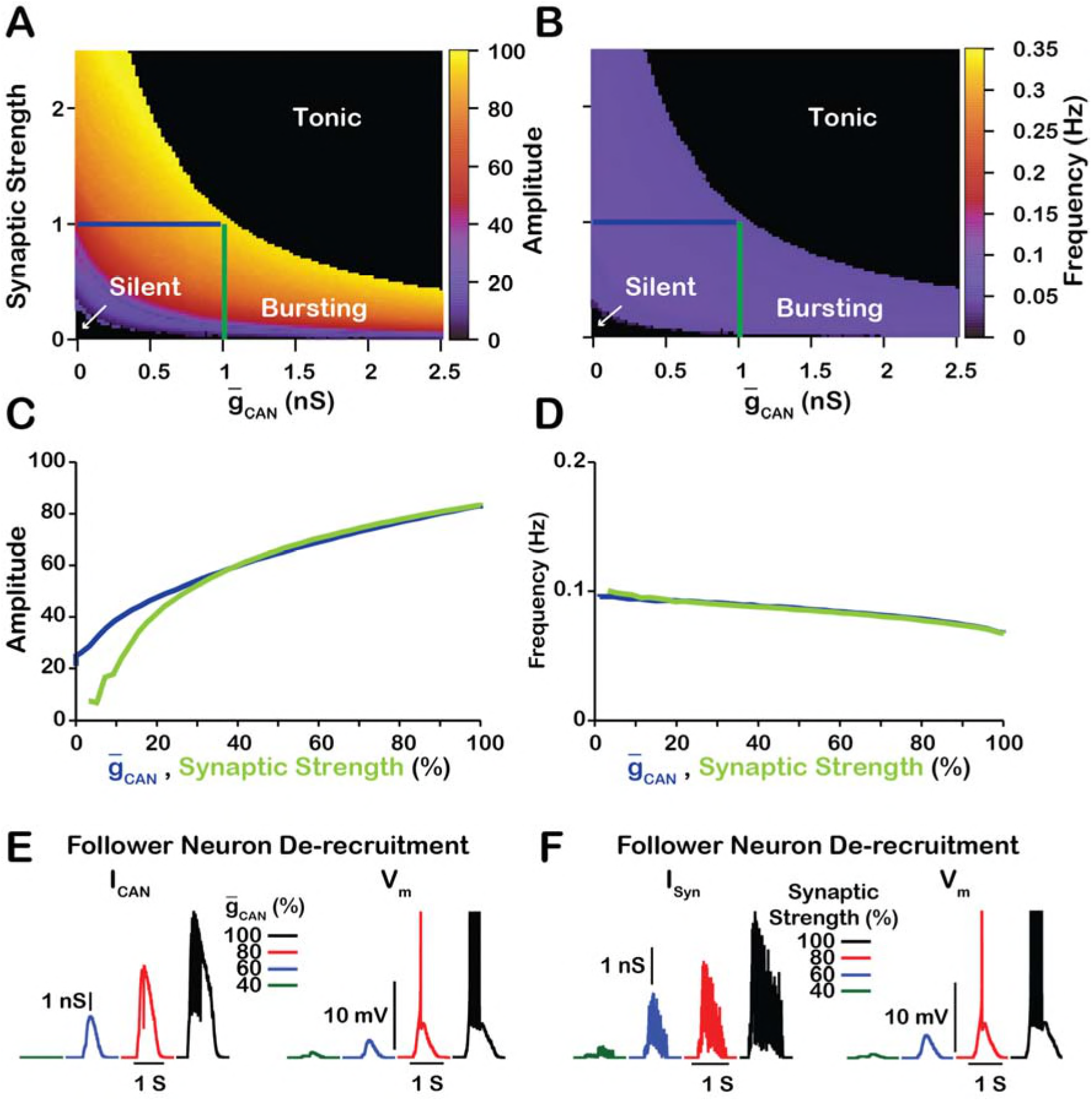
Manipulations of synaptic strength 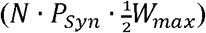 and 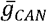 have equivalent effects on network activity amplitude, frequency and recruitment of follower neurons. **(A & B)** Relationship between 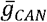, synaptic strength and the amplitude and frequency in the *Ca*_*Syn*_ network. Notice the symmetry about the X=Y line in panels **A** and **B,** which, indicates that changes in 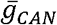 and or synaptic strength are qualitatively equivalent. Synaptic strength was changed by varying *W*_*max*_. **(C)** Relationship between network activity amplitude and the reduction of 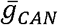 (BLUE) or synaptic strength (GREEN). **(D)** Relationship between network frequency and the reduction of 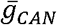 (BLUE) or synaptic strength (GREEN). **(E & F)** Decreasing 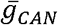 or synaptic strength de-recruits follower neurons by reducing the inspiratory drive potential, indicated by the amplitude of subthreshold depolarization, right traces. The solid blue and green lines in panels **A** and **B** represent the location in the 2D parameter space of the corresponding blue and green curves in **C** and **D**. The action potentials in the right traces of **E** and **F** are truncated to show the change in neuronal inspiratory drive potential. The parameters used for these simulations are *Ca*_*Syn*_: 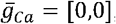, *P*_*Ca*_*= 0.01*, *P*_*Syn*_*= 1.0* and *W*_*max*_*= var*

### Robustness of Amplitude and Frequency Effects

We also examined if the effects are conserved in both the *Ca*_*V*_ and *Ca*_*Syn*_ networks over a range of network parameters. To test this, we investigated the dependence of network activity amplitude and frequency on 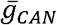 and average synaptic weight for *Ca*_*Syn*_ and *Ca*_*V*_ networks with high (*P*_*Syn*_*= 1*) and low (*P*_*Syn*_ = 0.05) connection probabilities, and high (*g*_*Ca*_*= 0.1 ns, P*_*Ca*_*= 0.1*), medium (*g*_*Ca*_*= 0.01 ns, P*_*Ca*_*= 0.01*) and low (*g*_*Ca*_*= 0.001 ns, P*_*Ca*_*= 0.005*) strengths of calcium sources (Figures 5 and 6). We found that changing the synaptic connection probability and changing the strength of the calcium sources has no effect on the general relationship between 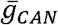 and the amplitude or frequency of bursts in the *Ca*_*V*_ or *Ca*_*Syn*_ networks. In other words, the general effect of increasing 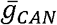 on amplitude and frequency is conserved in both networks regardless of the synaptic connection probability or strength of the calcium sources. Increasing the strength of the calcium sources does, however, affects the range of possible 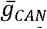 values where both networks produce rhythmic activity.

**Figure 5.**
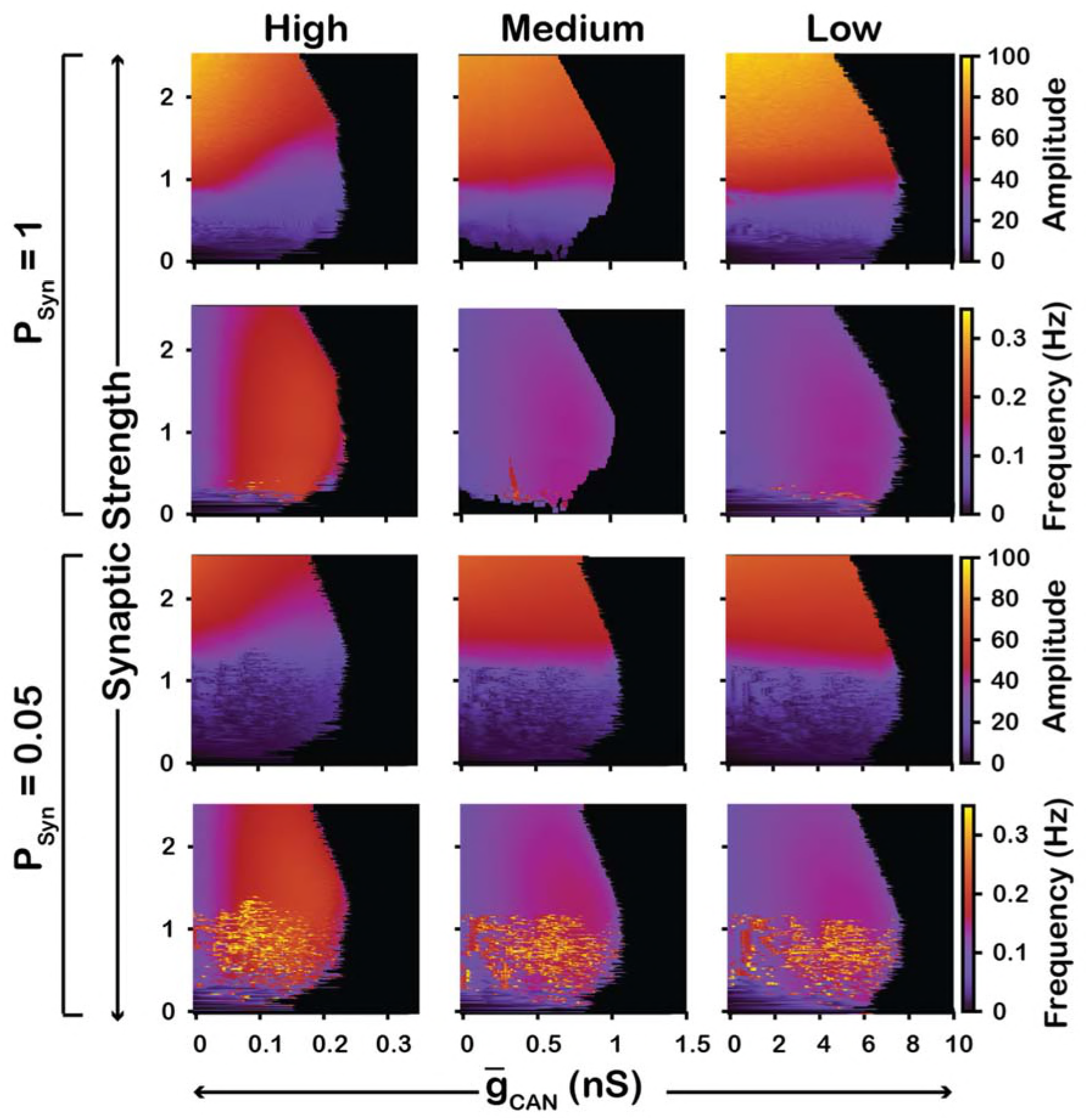
Robustness of amplitude and frequency effects to changes in 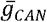 and synaptic strength in the *Ca*_*V*_ network for ‘high’(left), ‘medium’ (middle) and ‘low’ (right) conductance of the voltage-gated calcium channel *I*_*Ca*_ as well as ‘high’(top) and ‘low’ (bottom) network connection probabilities. Amplitude and frequency are indicated by color (scale bar at right). Black regions indicate tonic network activity.

**Figure 6.**
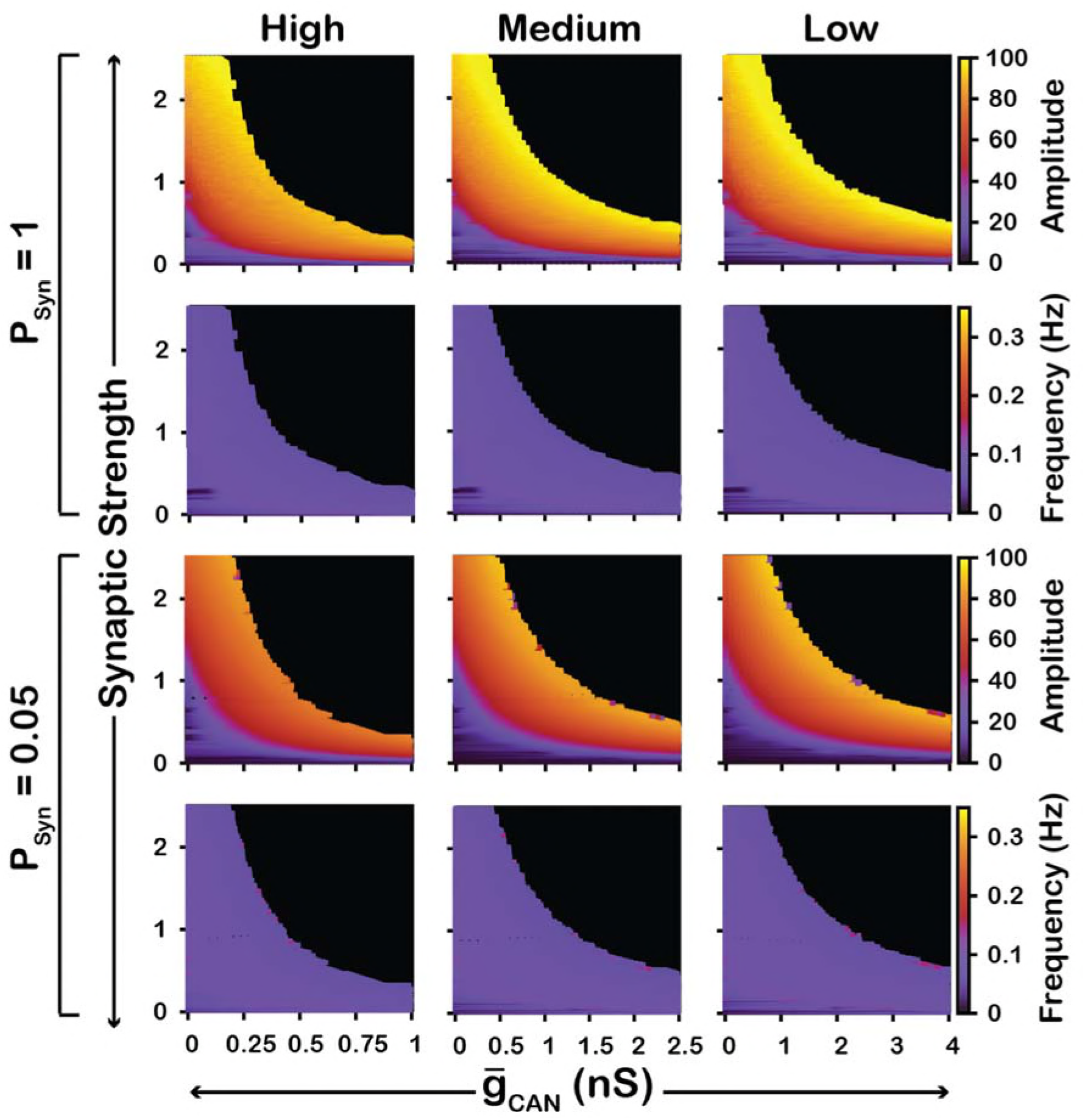
Robustness of amplitude and frequency effects to changes in 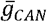 and synaptic strength in the *Ca*_*Syn*_ network for ‘high’(left), ‘medium’ (middle) and ‘low’ (right) calcium conductance in synaptic currents as well as ‘high’(top) and ‘low’ (bottom) network connection probabilities. Amplitude and frequency are indicated by color (scale bar at right). Black regions indicate tonic network activity.

To summarize, in the *Ca*_*V*_ model, increasing 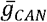 increases frequency, through increased excitability but has no effect on amplitude. In contrast, in the *Ca*_*Syn*_ model, increasing 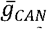 slightly decreases frequency and increases amplitude. In this case, increasing 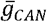 acts as a mechanism to increase the inspiratory drive potential and recruit previously silent neurons. Additionally, these features of the *Ca*_*V*_ and *Ca*_*Syn*_ models are robust and conserved across a wide range of network parameters.

### Intracellular Calcium Transients Activating *I*_*CAN*_ Primarily Result from Synaptically-Activated Sources

In experiments where *I*_*CAN*_ was blocked by bath application of FFA or 9-phenanthrol (Koizumi et al., 2018), the amplitude of network oscillations was strongly reduced and their frequency remained unchanged. Our model revealed that the effects of *I*_*CAN*_ blockade on amplitude and frequency depend on the source(s) of intracellular calcium (see Figs. 1 and 2). If the calcium influx is exclusively voltage-gated, our model predicts that *I*_*CAN*_ blockade will have no effect on amplitude but reduce the frequency. In contrast, if the calcium source is exclusively synaptically gated, our model predicts that blocking *I*_*CAN*_ will strongly reduce the amplitude and slightly increase the frequency. Therefore, a multi-fold decrease in amplitude, seen experimentally, is consistent with the synaptically driven calcium influx mechanisms, while constant bursting frequency may be due to calcium influx through both voltage-and synaptically gated channels. Following predictions above, to reproduce experimental data, we incorporated both mechanisms in the model and inferred their individual contributions by finding the best fit. We found that the best match is observed (Fig. 7) if synaptically mediated and voltage gated calcium influxes comprise about 95% and 5% of the total calcium influx, respectively.

**Figure 7.**
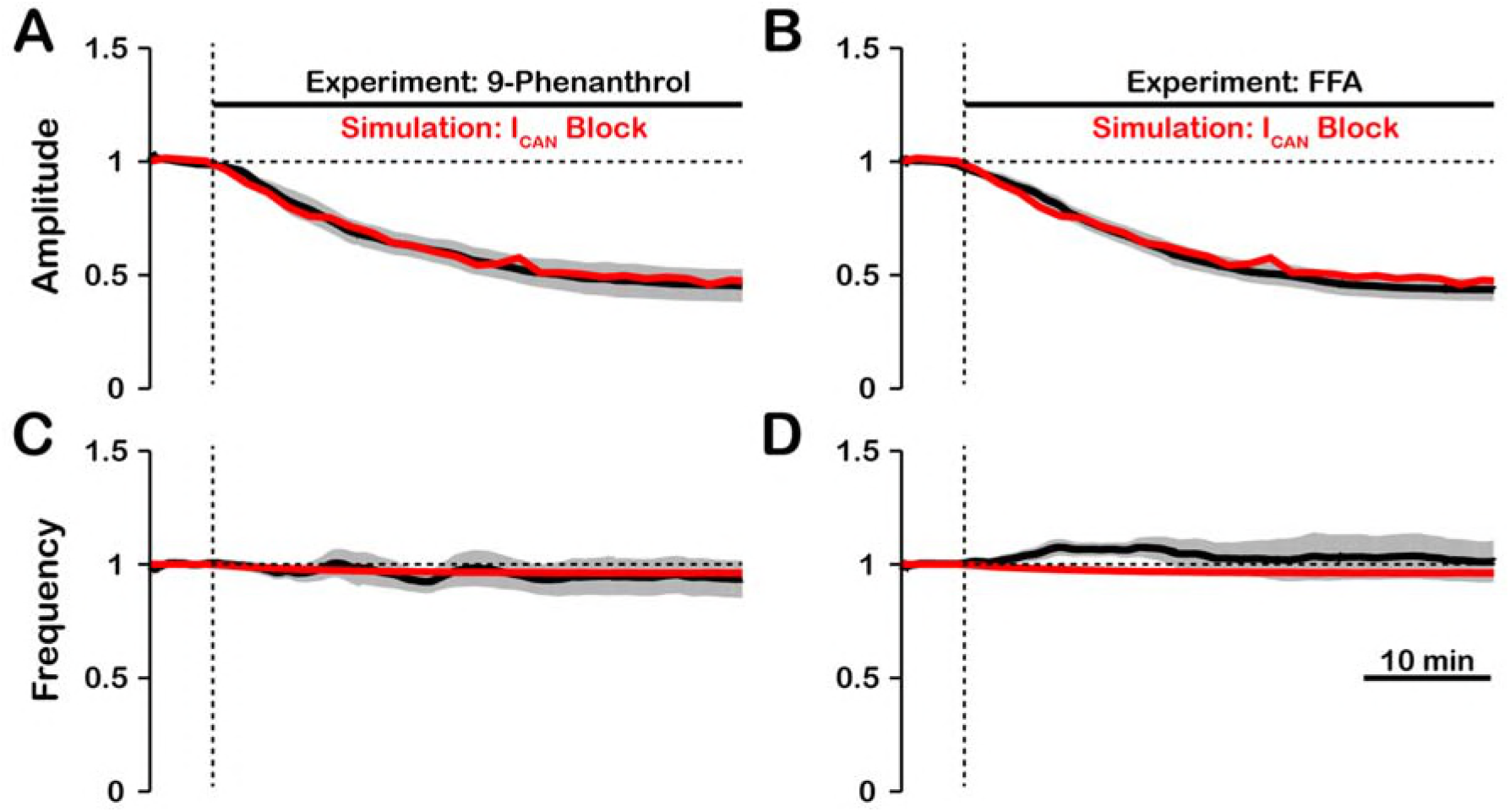
Experimental and simulated pharmacological blockade of *I*_*CAN*_ by (**A & C**) 9-phenanthrol and (**B & D**) flufenamic acid (FFA). Both voltage-gated and synaptic sources of intracellular calcium are included. Experimental blockade of *I*_*CAN*_ (black) by 9-phenanthrol and FFA significantly reduce the (**A & B**) amplitude of network oscillations while having little effect on (**C & D**) frequency. The black line represents the mean and the gray is the S.E.M. of experimental ?XII output recorded from neonatal rat brainstem slices in vitro, reproduced from Koizumi et al., 2018. Simulated blockade of *I*_*CAN*_ (red) closely matches the reduction in (**A & B**) amplitude of network oscillations and slight decrease in (**C & D**) frequency seen with 9-phenanthrol and FFA. Simulated and experimental blockade begins at the vertical dashed line. Blockade was simulated by exponential decay of 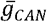 with the following parameters: 9-phenanthrol: *γ*_*Block*_*= 0.85*, *τ*_*Block*_*= 357s*; FFA: *γ*_*Block*_*= 0.92*, *τ*_*Block*_*= 415s*. The network parameters are: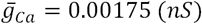, *P*_*Ca*_*= 0.0275*, *P*_*Syn*_*= 0.05* and *W*_*max*_*= 0.096 (ns)*.

### *I*_*NaP*_-dependent and [*Ca*]_*i*_-*I*_*CAN*_ Sensitive Intrinsic Bursting

In our model, we included *I*_*NaP*_, *I*_*CAN*_ as well as voltage-gated and synaptic mechanisms of *Ca*^*2+*^ influx. Activation of *I*_*CAN*_ by *Ca*_*Syn*_ is the equivalent mechanism used in other computational group-pacemaker models (Rubin et al., 2009; Song et al., 2015). Burst generation and termination in our model, however, are dependent on *I*_*NaP*_ (Butera et al., 1999a). We investigated the sensitivity of intrinsic bursting in our model to *I*_*NaP*_ and calcium channel blockade (Fig. 8). Intrinsic bursting was identified in neurons by zeroing the synaptic weights to simulate synaptic blockade. *I*_*NaP*_ and *I*_*Ca*_ blockade was simulated by setting 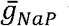 and 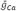 to *0*. We found that after decoupling the network (*W*_*max*_*= 0*) a subset of neurons remained rhythmically active (7%) and that these were all neurons with a high *I*_*NaP*_ conductance. In these rhythmically active neurons, bursting was abolished in all neurons by *I*_*NaP*_ blockade. Interestingly, *I*_*Ca*_ blockade applied before *I*_*NaP*_ blockade abolished intrinsic bursting in 2 of the 7 neurons and *I*_*NaP*_ blockade applied afterwards abolished intrinsic bursting in the remaining 5 neurons. Although only one rhythmogenic (*I*_*NaP*_-based) mechanism exists in this model, bursting in a subset of these neurons is calcium sensitive. In calcium-sensitive bursters, Ca^2+^ blockade abolishes bursting by reducing the intracellular calcium concentration and, hence, *I*_*CAN*_ activation, which ultimately reduces excitability.

**Figure 8.**
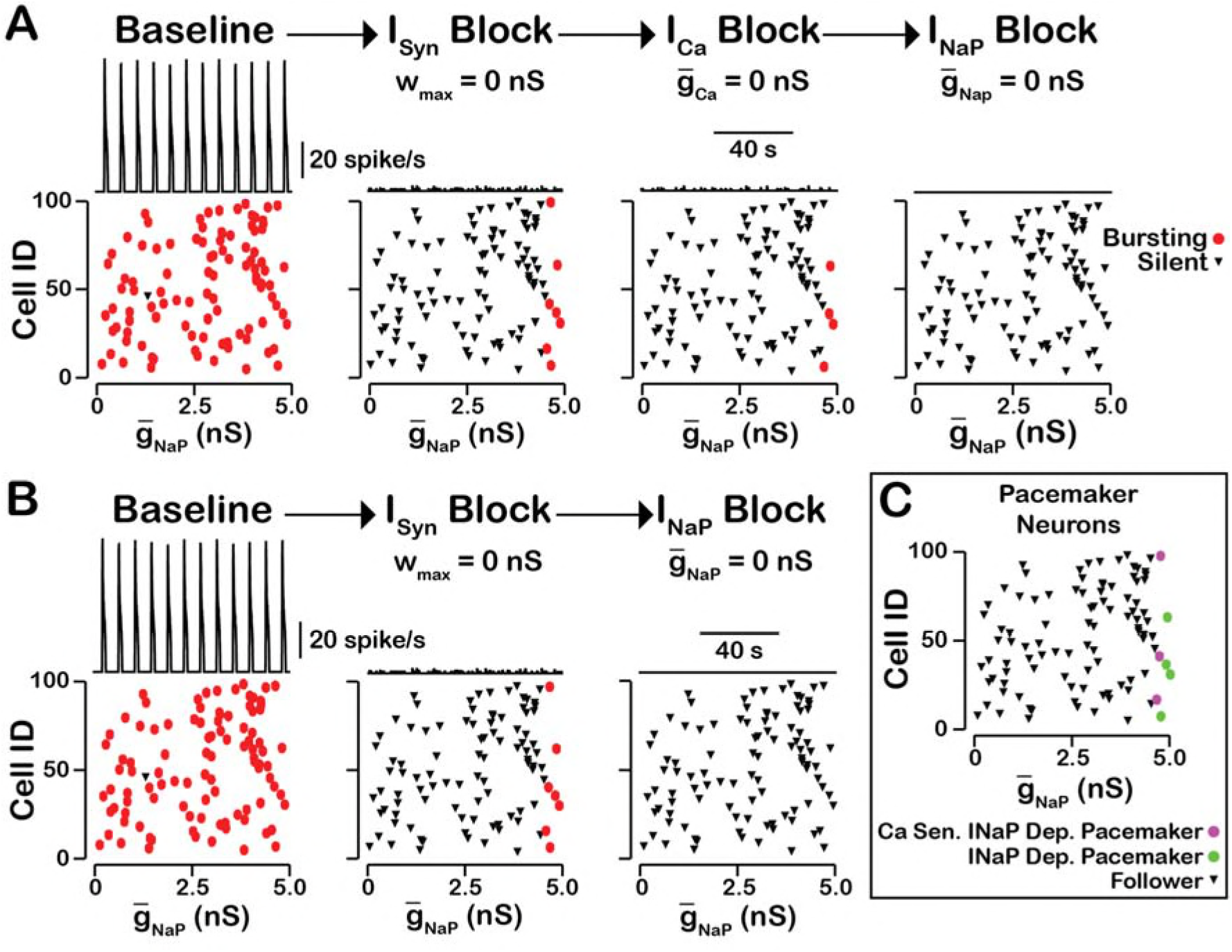
*I*_*NaP*_-dependent and *Ca2+* sensitive intrinsic bursting. **(A)** From left to right, intrinsic bursters are first identified by blocking synaptic connections. Then, calcium sensitive neurons are silenced and identified by *I*_*Ca*_ blockade. The remaining neurons are identifed as sensitive to *I*_*NaP*_ block. Top traces show the network output and Cell ID vs. 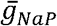 scatter plots identify silent and bursting neurons under each condition. **(B)** *I*_*NaP*_ blockade after synaptic blockade eleminates bursting in all neurons. Therefore, all intrinsic bursters are *I*_*NaP*_ dependent. **(C)** Identification of calcium sensitive and *I*_*NaP*_-dependent as well as calcium insensitive and *I*_*NaP*_-dependent intrinsic bursters. Notice that only the neurons with the highest value of 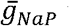 are intrinsic bursters and that a subset of these neurons are sensitive to calcium blockade but all are dependent on *I*_*NaP*_. The network parameters are: 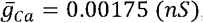, *P*_*Ca*_*= 0.0275*, *P*_*Syn*_*= 0.05* and *W*_*max*_*= 0.096 (ns)*.

### The Rhythmogenic Kernel

Our simulations have shown that the primary role of *I*_*CAN*_ is amplitude but not oscillation frequency modulation with little or no effect on network activity frequency. Here we examined the neurons that remain active and maintain rhythm after *I*_*CAN*_ blockade (Fig. 9). We found that the neurons that remain active are primarily neurons with the highest 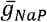 and that bursting in these neurons is dependent on *I*_*NaP*_. Some variability exists and neurons with relatively low 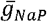 value can remain active due to synaptic interactions while a neuron with a slightly higher 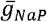 without sufficient synaptic input may become silent. These neurons, that remain active after compete blockade of *I*_*CAN*_, form a *I*_*NaP*_-dependent kernel of a rhythm generating circuit.

**Figure 9.**
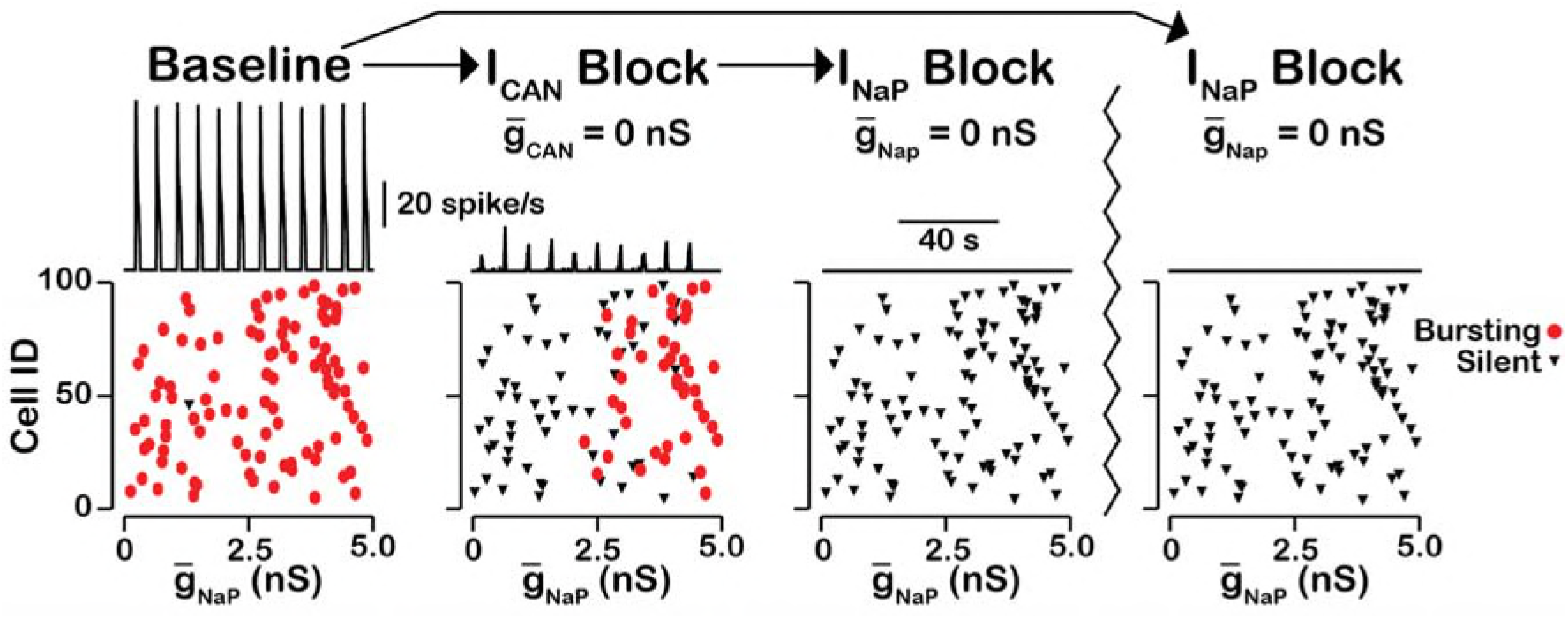
*I*_*CAN*_ blockade reveals an *I*_*NaP*_-dependent rhythmogenic kernel. The top traces show the network output at baseline, after *I*_*CAN*_ blockade and *I*_*NaP*_ blockade. The bottom Cell ID vs. 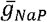 scatter plots identify silent and bursting neurons in each conditon. Notice that only neurons with relitively high 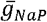 remain active after *I*_*CAN*_ block. The network parameters used are: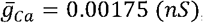, *P*_*Ca*_*= 0.0275*, *P*_*Syn*_*= 0.05* and *W*_*max*_*= 0.096 (ns)*.

### Network Activity Amplitude and Neuronal Spiking Frequency Correlate with Intracellular Calcium Transients

Dynamic calcium imaging has been utilized to assess activity of individual pre-BötC excitatory neurons and populations of these excitatory neurons in vitro during pharmacological inhibition of *I*_*CAN*_/TRPM4 (Koizumi et al., 2018). In such imaging-based analyses of excitatory network activity, the sources of calcium transients are not precisely known, but the calcium transients are assumed to correlate with the spiking frequency of neurons and provide measurements that are correlated with circuit activity as a whole, as the experimental data suggest. We compared the network activity characterized by the average intracellular calcium concentration and the network firing. We found that for a single network burst, the average intracellular calcium concentration and network firing are highly correlated (Fig. 10A).

**Figure 10.**
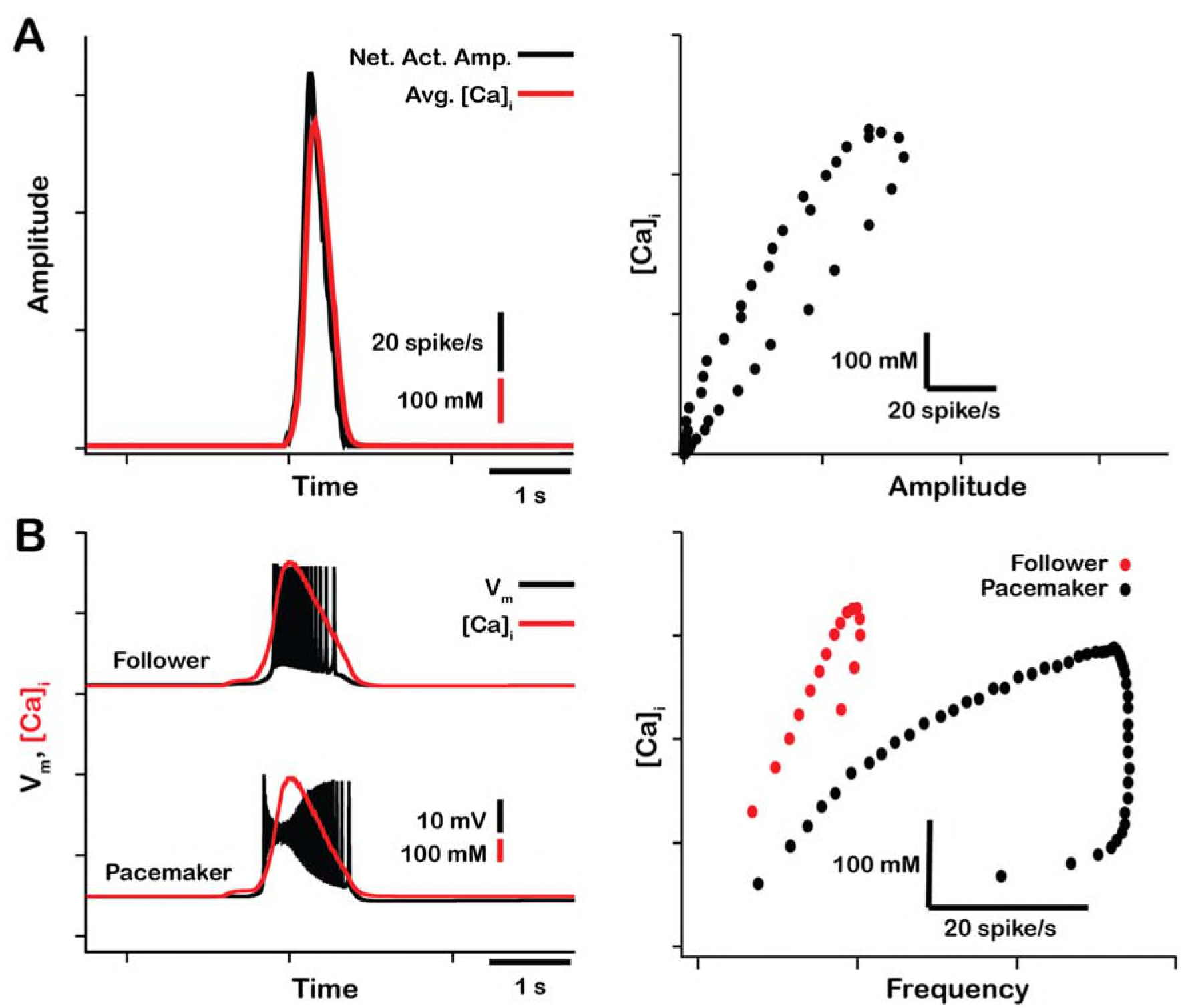
Correlations between network and neuronal spiking activity and intracellular calcium influx. **A)** Comparison of network activity amplitude and the average intracellular calcium concentration across a single network burst (left). Network activity amplitude and the average network intracellular calcium concentration are highly correlated (right). **B)** Comparison of spiking frequency and intracellular calcium concentration for a single burst in a typical follower and pacemaker neuron (left). Spiking frequency is highly correlated with the intracellular calcium transient in follower neurons (right, red). In the pacemaker neuron calcium influx occurs after spiking is initiated, consequently the correlation between spiking frequency and the intracellular calcium concentration is poor at the beginning of the bursts (right, black). The network parameters used are: 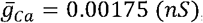, *P*_*Ca*_ = 0.0275, *P*_*Syn*_ = 0.05 and *W*_*max*_ = 0.096 (*ns*).

We also compared intracellular calcium transients and spiking frequency in individual pacemaker and follower neurons (Fig. 10B). Pacemaker neurons drive network oscillations and begin firing before the rest of the network. As a result, in the pacemaker neurons the action-potential generation precedes calcium influx. Consequently, in pacemaker neurons the [*Ca*]_*i*_ and firing rate correlate poorly in the first portion of the burst. In follower neurons bursting is dependent on synaptic input and recruitment through *I*_*CAN*_ activation. Thus, in follower neurons, which make up most neurons, the [*Ca*]_*i*_ and action potential firing rate are highly correlated.

### Low Network Connection Probability Increases the Variability of the Δ[*Ca*]_*i*_ in Individual Neurons During *I*_*CAN*_ Blockade

In our model, synaptically mediated calcium influx into the cell is proportional to the total synaptic current through its membrane. The synaptic current is determined by the number of synapses, the strength of the connections (synaptic weights) and the firing rates of the pre-synaptic neurons. The synaptic connection probability (*P*_*Syn*_) together with the total number of neurons in the network determines the average number of connections each neuron receives. We examined the relationship between connection probability and the change in [*Ca*]_*i*_ during simulated *I*_*CAN*_ blockade (Fig. 11). We found that the *P*_*Syn*_ has no effect on the relationship between amplitude, frequency or calcium transients at the network level provided that the synaptic strength remains constant 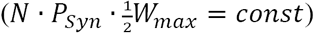 (Fig. 11A, B). Additionally, regardless of *P*_*Syn*_, the network activity amplitude and average intracellular calcium concentration are highly correlated. *P*_*Syn*_ does however affects the change in the peak [*Ca*]_*i*_ in individual neurons. In a network with a high connection probability (*P*_*Syn*_ = 1) the synaptic current/calcium transient is nearly identical for all neurons and therefore the change in [*Ca*]_*i*_ during *I*_*CAN*_ blockade is approximately the same for each neuron (Fig. 11C). In a sparsely connected network the synaptic current and calcium influx are more variable and reflect the heterogeneity in spiking frequency of the pre-synaptic neurons (Fig. 11D). Interestingly, in a network with low connection probability (*P*_*Syn*_*< 0.1*), the peak [*Ca*]_*i*_ transient in some neurons increases when *I*_*CAN*_ is blocked (Fig. 11E).

**Figure 11.**
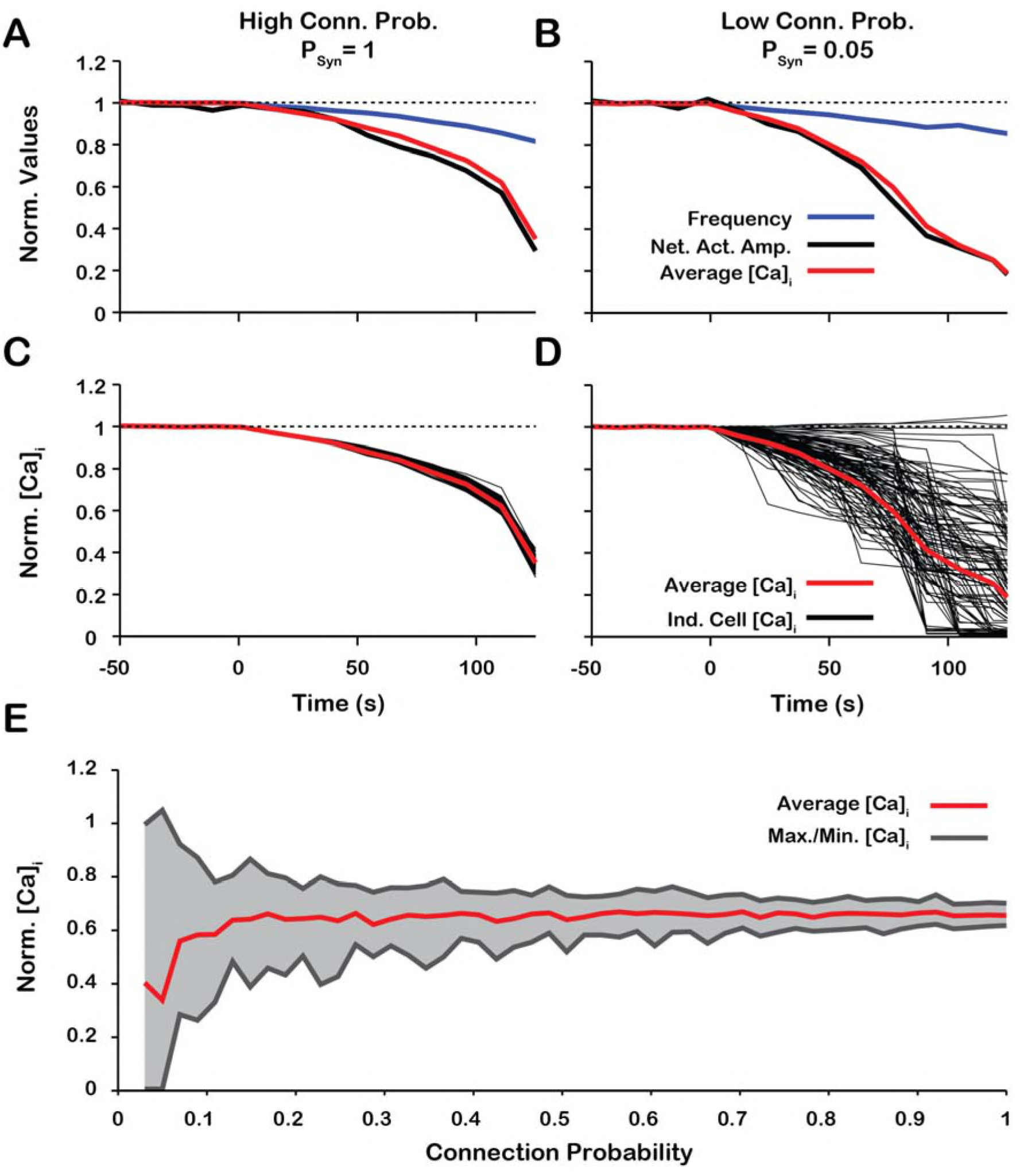
Low network connection probability increases the variability of changes in the peak [*Ca*]_*i*_ in individual neurons during *I*_*CAN*_ blockade. (**A** & **B**) Effect of *I*_*CAN*_ blockade on network activity amplitude, network calcium amplitude and frequency for network connection probabilities **A)** P = 1 and **B)** P = 0.05. (**C** & **D**) Effect of *I*_*CAN*_ blockade on changes in the magnitude of peak cellular calcium transients for network connection probabilities C) *P*_*Syn*_*= 1* and D) *P*_*Syn*_*= 0.05*. E) Maximum, minimum and average change in the peak intracellular calcium transient of individual neurons as a function of synaptic connection probability. All curves in A through E are normalized to their baseline values. Synaptic weight was adjusted to keep the average synaptic strength 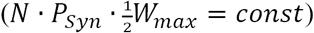 constant. Notice that lowering the synaptic connection probability increases the variability in the peak intracellular calcium concentration during *I*_*CAN*_ blockade. Interestingly, for connection probabilities below approximately 5%, blocking 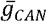 can increase the peak calcium transient in a small subset of neurons. The network parameters used are: 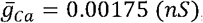 and *P*_*Ca*_ = 0.0275 and *W*_*max*_*= var*.

## 4. Discussion

Establishing cellular and circuit mechanisms generating the rhythm and amplitude of respiratory oscillations in the mammalian brainstem pre-BötC has remained an unsolved problem of wide-spread interest in neurophysiology since this structure, essential for breathing to support mammalian life, was discovered nearly three decades ago (Smith et al., 1991). Our objective in this theoretical study was to re-examine and further define contributions of two of the main currently proposed neuronal biophysical mechanisms operating in pre-BötC excitatory circuits, specifically mechanisms involving *I*_*CAN*_ activated by neuronal calcium fluxes and voltage-dependent *I*_*NaP*_ in the circuit neurons. While these sodium-and calcium-based mechanisms have been studied extensively over the past two-decades and shown experimentally to be integrated in pre-BötC circuits, their actual roles in circuit operation are continuously debated and unresolved. Both mechanisms have been proposed to be fundamentally involved in rhythm generation either separately or in combination, as plausibly shown from previous theoretical modeling studies. Furthermore, the process of rhythm generation in pre-BötC circuits must be associated with an amplitude of excitatory circuit activity sufficient to drive downstream circuits to produce adequate respiratory motor output. Biophysical mechanisms involved in generating excitatory population activity amplitude have also not been established. Our analysis is motivated by the recent experimental observations obtained from neonatal rodent slices isolating pre-BötC circuits in vitro that inhibition of the endogenously active *I*_*CAN*_/TRPM4 strongly reduces the amplitude of network oscillations within pre-BötC circuits but have little effect on oscillation frequency (Koizumi et al., 2018). This is contrary to the proposed *I*_*CAN*_-based models for rhythm generation in the isolated pre-BötC and indicates a fundamentally different functional organization of pre-BötC circuits, in terms of oscillatory frequency and amplitude generation, that needs to be defined.

We accordingly analyzed the role of *I*_*CAN*_ and possible sources of intracellular calcium transients activating this conductance and found that the effect of simulated *I*_*CAN*_ blockade on amplitude and frequency is highly dependent on the source(s) of intracellular calcium, which is also a central issue to be resolved. In the case where *Ca*_*Syn*_ is the primary intracellular calcium source, *I*_*CAN*_ blockade generates a large reduction in network activity amplitude. In contrast, when *Ca*_*V*_ is the only intracellular calcium source, *I*_*CAN*_ blockade has little effect on network activity amplitude and primarily affects the population bursting frequency that is caused by decreased excitability. Additionally, we show that activation of *I*_*CAN*_ by *Ca*_*Syn*_ functions as a mechanism to augment the inspiratory drive potential and amplitude of population activity, and that this effect is similar to increasing the synaptic coupling strength within the network. Therefore, in the case of *Ca*_*Syn*_, blockade of *I*_*CAN*_ reduces the inspiratory drive potential causing de-recruitment of non-rhythmogenic follower neurons and reduction of network activity amplitude. In a model where *I*_*CAN*_ is activated by both *Ca*_*V*_ and *Ca*_*Syn*_ with contributions of 5% and 95% respectively, we show that simulated blockade of *I*_*CAN*_ generates a large reduction in network population activity amplitude and a slight decrease in frequency. This closely reproduces experimental blockade of *I*_*CAN*_/TRPM4 by either 9-phenanthrol or FFA (Fig. 7). Finally, we showed that the change in the peak calcium transients for individual neurons during *I*_*CAN*_ blockade match experimental data particularly at relatively low connection probabilities (*P*_*Syn*_∼ < 0.1).

### Role of *I*_*CAN*_ in the pre-BötC Respiratory Network

The hypothesis that *I*_*CAN*_ is involved in generation of the inspiratory rhythm is based on experimental observations from in vitro mouse medullary slice preparations (Pena et al., 2004; Thoby-Brisson & Ramirez, 2001), and in silico modeling studies (Jasinski et al., 2013; Rubin, Hayes, et al., 2009; Toporikova & Butera, 2011). Theories of *I*_*CAN*_-dependent bursting rely on intracellular *Ca*^*2+*^ signaling mechanisms that have not been well defined.

Two models of *I*_*CAN*_-dependent rhythmic bursting in vitro have been proposed and are referred to as the “dual pacemaker” and “group pacemaker” models. In the dual pacemaker model, two types of pacemaker neurons are proposed that are either *I*_*NaP*_-dependent (riluzole sensitive) or *I*_*CAN*_-dependent (*Cd*^*2+*^ sensitive) intrinsic bursters (see Rybak et al., 2014 for review). In this model network, oscillations are thought to originate from these pacemaker neurons which through excitatory synaptic interactions synchronize bursting and drive activity of follower neurons within the pre-BötC. Although pacemaker neurons sensitive to neuronal *Ca*^*2+*^ flux blockade through *Ca*_*V*_ have been reported (Pena et al., 2004; Thoby-Brisson & Ramirez, 2001), the source and mechanism driving intracellular *Ca*^*2+*^oscillations has not been described. Computational models of *I*_*CAN*_-dependent pacemaker neurons rely on controversial mechanisms for burst initiation and terminations, e.g. IP3-dependent *Ca*^*2+*^oscillations (Toporikova & Butera, 2011; Del Negro et al., 2010), that have been questioned from recent negative experimental results (Beltran-Parrazal et al., 2012; Toporikova et al., 2015).

In versions of the “group pacemaker” model (Rubin et al., 2009; Feldman & Del Negro, m2006; Del Negro et al., 2010) network oscillations are initiated through recurrent synaptic excitation that trigger postsynaptic *Ca*^*2+*^influx. Subsequent *I*_*CAN*_ activation generates membrane depolarization (inspiratory drive potential) to drive neuronal bursting. Synaptically triggered *Ca*^*2+*^influx and the contribution of *I*_*CAN*_ to the inspiratory drive potential of individual pre-BötC neurons are experimentally supported (Mironov, 2008; Pace et al., 2007), however the mechanism of burst termination remains unclear. Again, the computational group-pacemaker models that have been explored (Rubin, et al., 2009) rely on as yet unproven mechanisms for burst termination, and in some cases lack key biophysical features of the pre-BötC neurons such as voltage-dependent frequency control and expression of *I*_*NaP*_.

In our model, we showed that blockade of either *I*_*CAN*_ or synaptic interactions produce qualitatively equivalent effects on network population activity amplitude and frequency when the calcium transients are primarily generated from synaptic sources (Fig.4). Consequently, our model predicts that blockade of *I*_*CAN*_ or synaptic interactions in the isolated pre-BötC in vitro will produce comparable effects on amplitude and frequency. This is the case as Johnson et al. (1994) showed that gradual blockade of synaptic interactions by low calcium solution significantly decreases network activity amplitude while having little effect of frequency, similar to the experiments where the *I*_*CAN*_ channel TRPM4 is blocked with 9-phenanthrol (Koizumi et al., 2018).

Overall, our new model simulations for the isolated pre-BötC excitatory network suggest that the role of *I*_*CAN*_/TRPM4 activation is to amplify excitatory synaptic drive in generating the amplitude of inspiratory population activity, independent of the biophysical mechanism generating inspiratory rhythm. We note that the recent experiments have also shown that in the more intact brainstem respiratory network that ordinarily generates patterns of inspiratory and expiratory activity, endogenous activation of *I*_*CAN*_/TRPM4 appears to augment the amplitude of both inspiratory and expiratory population activity, and hence these channels are fundamentally involved in inspiratory-expiratory pattern formation (Koizumi et al., 2018).

### Calcium Transients as Correlates of Activity

Neuronal calcium transients can arise from voltage-gated calcium sources, driven by action potentials, and serve as correlates of neuronal activity. We analyzed the correlation between calcium transients and inspiratory activity of individual inspiratory neurons as well as the entire network, particularly since dynamic calcium imaging has been utilized to assess activity of individual and populations of pre-BötC excitatory neurons in vitro during pharmacological inhibition of *I*_*CAN*_/TRPM4 (Koizumi et al., 2018). In our model, most of the calcium influx is synaptically-triggered and may occur within a given neuron in the absence of action potentials. We show that intracellular calcium transients are highly correlated with network and cellular activity. This is true across individual neuron bursts and when comparing changes in peak values of neuronal firing and intracellular calcium transients across the duration of an *I*_*CAN*_ blockade simulation. The correlation at the onset of bursting in pacemaker neurons are an exception. In these neurons, the correlations between the intracellular calcium concentration and the instantaneous firing rate across a single burst are not apparent at the onset of this burst. This is because pacemaker neurons start spiking before the rest of the network, which precedes synaptically triggered calcium influx.

Additionally, we examined the relative change in the peak calcium transients in single neurons as a function of *I*_*CAN*_ conductance. We show that in a subset of neurons the peak calcium transient increases with reduced *I*_*CAN*_. This result is surprising but is supported by the recent calcium imaging data (Koizumi et al., 2018). This occurs in neurons that receive most of their synaptic input from pacemaker neurons and our analyses suggest this is possible in sparse networks, i.e. with relatively low connection probability. In pacemaker neurons, *I*_*CAN*_ blockade leads to a reduction of their excitability resulting in an increased value of *I*_*NaP*_ inactivation gating variable at the burst onset. Thus, during the burst, the peak action potential frequency and the synaptic output from these neurons is increased with *I*_*CAN*_ blockade. Consequently, neurons that receive synaptic input from pacemaker neurons will see an increase in their peak calcium transients. In most neurons, however, synaptic input is received primarily from follower neurons. Since *I*_*CAN*_ blockade de-recruits follower neurons, the synaptic input and subsequent calcium influx in most decreases. Therefore, our model predicts that in a sparse network, blocking *I*_*CAN*_ results in very diverse responses at the cellular level with overall tendency to reduce intracellular calcium transients such that the amplitude of these transients averaged over the entire population decreases during *I*_*CAN*_ blockade while their frequency is unchanged. This is consistent with the experimental calcium imaging data (Koizumi et al., 2018).

### Synaptic Calcium Sources

Our model suggests that calcium transients in the pre-BötC are coupled to excitatory synaptic input, i.e. pre-synaptic glutamate release and binding to post-synaptic glutamate receptors triggers calcium entry. The specific mechanisms behind this process is unclear, however it is likely dependent on specific types of ionotropic or metabotropic glutamate receptors.

There are three subtypes of ionotropic glutamate receptors, N-methyl-D-aspartate (NMDA), Kainate (KAR), and α-amino-3-hydroxy-5-methyl-4-isoxazolepropionic acid (AMPA), all of which are expressed in the pre-BötC (Paarmann, Frermann, Keller, & Hollmann, 2000) and have varying degrees of calcium permeability. NMDA and AMPA are unlikely candidates for direct involvement in synaptically mediated calcium influx in the pre-BötC. Pharmacological blockade of NMDA receptors has no significant effect on the amplitude or frequency of XII motor output (Morgado-Valle & Feldman, 2007; Pace et al., 2007) and AMPA receptors in the pre-BötC show high expression of the subunit GluR2, which renders the AMPA ion channel pore impermeable to Ca^2+^ (Paarmann et al., 2000). It is possible, however, that AMPA mediated depolarization may trigger calcium influx indirectly through the voltage-gated calcium channel activation on the post-synaptic terminal. The contribution of the latter to synaptically triggered calcium influx is likely small as pharmacological blockade of L-, N-and P/Q-type calcium channels have no significant effect on XII motor output from the pre-BötC (Morgado-Valle et al., 2008).

Calcium permeability through KAR receptors is dependent on subunit expression. The KAR subunit GluK3 is highly expressed in the pre-BötC (Paarmann et al., 2000) and is calcium permeable (Perrais et al., 2009) making it a possible candidate for synaptically mediated calcium entry. Furthermore, GluK3 is insensitive to tonic glutamate release and only activated by large glutamate transients (Perrais et al., 2009). Consequently, GluK3 may only be activated when receiving synaptic input from a bursting presynaptic neuron which would presumably generate large glutamate transients. The role of GluK3 in the pre-BötC has not been investigated.

Metabotropic glutamate receptors (mGluR) indirectly activate ion channels through G-protein mediated signaling cascades. Group 1 mGluRs which include mGluR1 and mGluR5 are typically located on post-synaptic terminals (Shigemoto et al., 1997) and activation of group 1 mGluRs is commonly associated with calcium influx through calcium permeable channels (Berg et al., 2007; Endoh, 2004; Mironov, 2008) and calcium release from intracellular calcium stores (Pace et al., 2007).

In the pre-BötC, mGluR1/5 are thought to contribute to calcium influx by triggering the release of calcium from intracellular stores (Pace et al., 2007) and/or the activation of the transient receptor potential C3 (TRPC3) channel (Ben-Mabrouk & Tryba, 2010). Blockade of mGluR1/5 reduces the inspiratory drive potential in pre-BötC neurons and reduces XII motor output (Pace et al., 2007) which is consistent with the effects of *I*_*CAN*_/TRPM4 blockade (Pace et al., 2007). TRPC3 is a calcium permeable channel (Thebault et al., 2005) that is associated with calcium signaling (Hartmann et al., 2011), store-operated calcium entry (Kwan et al., 2004), and synaptic transmission (Hartmann et al., 2011). TRPC3 is activated by diacylglycerol (DAG) (Clapham, 2003) which is formed after synaptic activation of mGluR1/5. TRPC3 is highly expressed in the pre-BötC and was hypothesized to underlie *I*_*CAN*_ activation in the pre-BötC (Ben-Mabrouk & Tryba, 2010) and other brain regions (Amaral & Pozzo-Miller, 2007; Zitt et al., 1997). Furthermore, TRPC3 and *I*_*CAN*_ have been shown to underlie slow excitatory post synaptic current (sEPSC) (Hartmann et al., 2008; Hartmann et al., 2011). This is consistent with our model since *I*_*CAN*_ activation is dependent on synaptically triggered calcium entry, and the calcium dynamics are slower than the fast AMPA based current *I*_*Syn*_. Therefore, in our model, *I*_*CAN*_ decays relatively slowly and, hence, can be treated as a sEPSC.

In the pre-BötC, the effect of TRPC3 blockade by 3-pyrazole on network amplitude is remarkably similar to blockade of TRPM4 (Koizumi et al., 2018). This suggests that the *I*_*CAN*_/TRPM4 activation may be dependent on/coupled to TRPC3. A possible explanation is that TRPC3 mediates synaptically-triggered calcium entry. It is also likely that TRPC3 plays a role in maintaining background calcium concentration levels. We tested this hypothesis by simulating the blockade of synaptically-triggered calcium influx while simultaneously lowering the background calcium concentration (Fig. 12). These simulations generated large reductions in activity amplitude with no effect on frequency which are consistent with data from experiments where TRPC3 is blocked using 3-pyrazole (Koizumi et al., 2018). This indirectly suggests that TRPC3 is critical for synaptically-triggered calcium entry and subsequent *I*_*CAN*_ activation.

**Figure 12.**
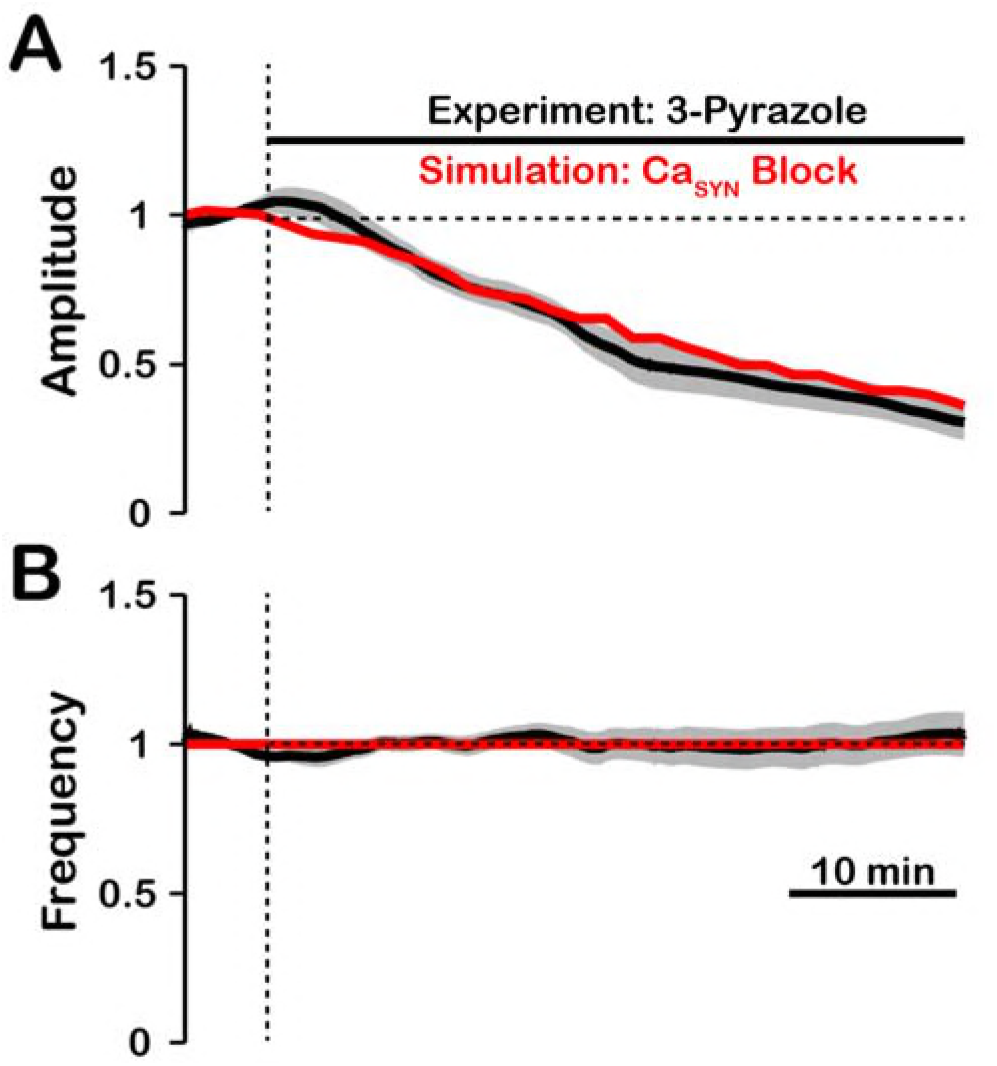
Comparison of experimental (black) and simulated (red) TRPC3 blockade (by *Ca*_*Syn*_ block) on network activity amplitude (**A**) and frequency (**B**). Simulated and experimental blockade begins at the vertical dashed line. The black line represents the mean and the gray band represents the mean S.E.M. of experimental ?XII output recorded from neonatal rat brainstem slices in vitro, reproduced from Koizumi et al., 2018. Blockade was simulated by exponential decay of *P*_*Ca*_ with the following parameters: 3-Pyrazole: *γ*_*Block*_ = 1.0, *τ*_*Block*_ = 522.5*s*. The network parameters are: 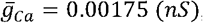, *P*_*Ca*_*= 0.0275*, *P*_*Syn*_*= 0.05* and *W*_*max*_*= 0.096 (ns)*.

### *I*_*NaP*_*-*dependent Rhythmogenic Kernel

*I*_*NaP*_ is a conductance present ubiquitously in pre-BötC inspiratory neurons, and is established to underlie intrinsic oscillatory neuronal bursting in the absence of excitatory synaptic interactions in neurons with sufficiently high *I*_*NaP*_ conductance densities (Koizumi & Smith, 2008; Koizumi et al., 2008). Accordingly, we randomly incorporated this conductance in our model excitatory neurons from a uniform statistical distribution to produce heterogeneity in *I*_*NaP*_ conductance density across the population. Our simulations indicate that the circuit neurons mostly with relatively high *I*_*NaP*_ conductance values underlie rhythm generation and remain active after compete blockade of *I*_*CAN*_ in our model network, thus forming a *I*_*NaP*_-dependent rhythmogenic kernel, including some neurons with intrinsic oscillatory bursting behavior when synaptically uncoupled.

Recently it has become apparent that there is functional heterogeneity within pre-BötC excitatory circuits, including distinct subpopulations of neurons involved in generating periodic sighs (Toporikova et al., 2015; Li et al., 2016), and the subpopulations generating regular inspiratory activity. Activity of both of these subpopulations in the pre-BötC isolated in vitro is proposed to be dependent on activation of *I*_*NaP*_ (Toporikova et al., 2015). Our experimental and modeling results suggest that within the normal inspiratory population, there are subpopulations distinguished by their role in rhythm versus amplitude generation due to biophysical properties: there is a *I*_*CAN*_/TRPM4-dependent recruitable population of excitatory neurons for burst amplitude generation and the *I*_*NaP*_-dependent rhythmogenic kernel population. The spatial arrangements of these two synaptically connected excitatory populations within the pre-BötC are currently unknown, and it remains an important experimental problem to identify the cells constituting the rhythmogenic kernel and their biophysical properties. This should now be possible, since our analysis and experimental results suggest that the rhythmically active neurons of the kernel population can be revealed and studied after pharmacologically inhibiting the *I*_*CAN*_/TRPM4-dependent inspiratory burst-generating population.

Recently a “burstlet theory” for emergent network rhythms has been proposed to account for inspiratory rhythm and pattern generation in the isolated pre-BötC in vitro (Kam et al., 2013; Del Negro et al., 2018). This theory postulates that a subpopulation of excitatory neurons generating small amplitude oscillations (burstlets) functions as the inspiratory rhythm generator that drives neurons that generate the larger amplitude, synchronized inspiratory population bursts. This concept emphasizes that subthreshold neuronal membrane oscillations need to be considered and that there is a neuronal subpopulation that functions to independently form the main inspiratory bursts. This is similar to our concept of distinct excitatory subpopulations generating the rhythm versus the amplitude of inspiratory oscillations. Biophysical mechanisms generating rhythmic burstlets and the large amplitude inspiratory population bursts in the burstlet theory are unknown. We have identified a major Ca^2+^-dependent conductance mechanism for inspiratory burst amplitude (pattern) generation and show theoretically how this mechanism may be coupled to excitatory synaptic interactions and is independent of the rhythm-generating mechanism. We also note that a basic property of *I*_*NaP*_ is its ability to generate subthreshold oscillations and promote burst synchronization (Butera et al., 1999b; Bacak et al., 2016). However, in contrast to our proposal for the mechanisms operating in the kernel rhythm-generating subpopulation, *I*_*NaP*_ with its favorable voltage-dependent and kinetic autorhythmic properties– is not proposed to be a basic biophysical mechanism for rhythm generation in the burstlet theory (Del Negro et al., 2018).

We emphasize that the above discussions regarding the role of *I*_*NaP*_ pertain to the isolated pre-BötC including in more mature rodent experimental preparations in situ where inspiratory rhythm generation has also been shown to be dependent on *I*_*NaP*_ (Smith et al., 2007). The analysis is more complex when the pre-BötC is embedded within interacting respiratory circuits in the intact nervous system generating the full complement of inspiratory and expiratory phase activity, where rhythmogenesis is tightly controlled by inhibitory circuit interactions and the contribution of *I*_*NaP*_ kinetic properties alone in setting the timing of inspiratory oscillations is diminished (Smith et al., 2007; Rubin et al., 2009; Richter & Smith, 2014).

### Conclusions

Based on our new data-driven computational model, distinct biophysical mechanisms are involved in generating the rhythm and amplitude of inspiratory oscillations in the isolated pre-BötC excitatory circuits. Inspiratory rhythm generation arises from a group of *I*_*NaP*_-dependent excitatory neurons, including cells with intrinsic oscillatory bursting properties, that form a rhythmogenic kernel. Rhythmic synaptic drive from these neurons triggers post-synaptic calcium transients, *I*_*CAN*_ activation, and subsequent membrane depolarization which drives bursting in the population of non-rhythmogenic follower neurons. We showed that activation of *I*_*CAN*_ by synaptically-driven calcium influx functions as a mechanism that amplifies the excitatory synaptic input to generate the inspiratory drive potential and population activity amplitude in these non-rhythmogenic neurons. Consequently, blockade of *I*_*CAN*_ causes a robust decrease in overall network activity amplitude via de-recruitment of these follower neurons without perturbations of the inspiratory rhythm, which is consistent with the results with experimental blockade of *I*_*CAN*_/TRPM4 channels. Our model provides a theoretical explanation for these recent paradigm-shifting experimental results that *I*_*CAN*_ is not fundamentally involved in generating the inspiratory rhythm and gives new insights into the functional operation of pre-BötC excitatory circuits.

## Materials and Methods

### Model Description

The model describes a network of *N* = 100 synaptically coupled excitatory neurons. Simulated neurons are comprised of a single compartment described using a Hodgkin Huxley formalism. For each neuron, the membrane potential *V*_m_ is given by the following current balance equation:

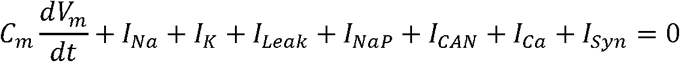

where *C*_*m*_ is the membrane capacitance, *I*_*Na*_, *I*_*K*_, *I*_*Leak*_, *I*_*NaP*_, *I*_*CAN*_, *I*_*Ca*_ and *I*_*Syn*_ are ionic currents through sodium, potassium, leak, persistent sodium, calcium activated non-selective cation, voltage-gated calcium, and synaptic channels, respectively. Description of these currents, synaptic interactions, and parameter values are taken from (Jasinski et al., 2013). The channel currents are defined as follows:

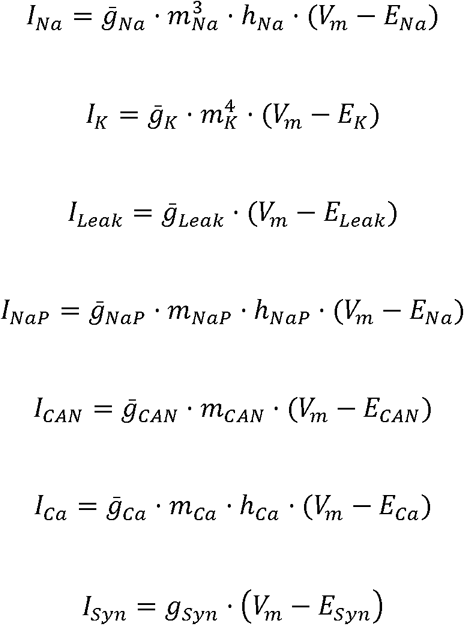

where 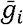 is the maximum conductance, *E*_*i*_ is the reversal potential, *m*_*i*_ and *h*_*i*_ are voltage dependent gating variables for channel activation and inactivation, respectively, and *i ∈ {Na, K, Leak, NaP, CAN, Ca, syn}*. The parameters 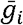 and *E*_*i*_ are given in Table 1.

**Table 1.** Model parameter values. The channel kinetics, intracellular Ca^2+^ dynamics and the corresponding parameter values, were derived from previous models (see (Jasinski et al., 2013) and the references therein).

| Channel | Parameters |
| --- | --- |
| $I_{Na}$ | $\bar{g}_{Na} = 150.0 \text{ nS}$, $E_{Na} = 55.0 \text{ mV}$,<br>$V_{m_{1/2}} = -43.8 \text{ mV}$, $k_m = 6.0 \text{ mV}$,<br>$V_{\tau_{m_{1/2}}} = -43.8 \text{ mV}$, $k_{\tau_m} = 14.0 \text{ mV}$, $\tau_{m_{max}} = 0.25 \text{ ms}$,<br>$V_{h_{1/2}} = -67.5 \text{ mV}$, $k_h = -10.8 \text{ mV}$,<br>$V_{\tau_{h_{1/2}}} = -67.5 \text{ mV}$, $k_{\tau_h} = 12.8 \text{ mV}$, $\tau_{h_{max}} = 8.46 \text{ ms}$ |
| $I_K$ | $\bar{g}_K = 160.0 \text{ nS}$, $E_K = -94.0 \text{ mV}$,<br>$A_\alpha = 0.01$, $B_\alpha = 44.0 \text{ mV}$, $\kappa_\alpha = 5.0 \text{ mV}$<br>$A_\beta = 0.17$, $B_\beta = 49.0 \text{ mV}$, $\kappa_\beta = 40.0 \text{ mV}$ |
| $I_{Leak}$ | $\bar{g}_{Leak} = 2.5 \text{ nS}$, $E_{Leak} = -68.0 \text{ mV}$, |
| $I_{NaP}$ | $\bar{g}_{NaP} \in [0.0, 5.0] \text{ nS}$,<br>$V_{m_{1/2}} = -47.1 \text{ mV}$, $k_m = 3.1 \text{ mV}$,<br>$V_{\tau_{m_{1/2}}} = -47.1 \text{ mV}$, $k_{\tau_m} = 6.2 \text{ mV}$, $\tau_{m_{max}} = 1.0 \text{ ms}$,<br>$V_{h_{1/2}} = -60.0 \text{ mV}$, $k_h = -9.0 \text{ mV}$,<br>$V_{\tau_{h_{1/2}}} = -60.0 \text{ mV}$, $k_{\tau_h} = 9.0 \text{ mV}$, $\tau_{h_{max}} = 5000 \text{ ms}$ |
| $I_{CAN}$ | $\bar{g}_{CAN} = 1.0 \text{ nS}, E_{CAN} = 0.0 \text{ mV},$<br>$Ca_{1/2} = 0.00074 \text{ mM}, n = 0.97$ |
| $I_{Ca}$ | $\bar{g}_{Ca} = 0.01 \text{ nS}, E_{Ca} = R \cdot T / F \cdot \ln([Ca]_{out} / [Ca]_{in}),$<br>$R = 8.314 \text{ J} / (\text{mol} \cdot \text{K}), T = 308.0 \text{ K},$<br>$F = 96.485 \text{ kC} / \text{mol}, [Ca]_{out} = 4.0 \text{ mM}$<br>$V_{m_{1/2}} = -27.5 \text{ mV}, k_m = 5.7 \text{ mV}, \tau_m = 0.5 \text{ ms},$<br>$V_{h_{1/2}} = -52.4 \text{ mV}, k_h = -5.2 \text{ mV}, \tau_h = 18.0 \text{ ms}$ |
| $Ca_{in}$ | $\alpha_{Ca} = 2.5 \cdot 10^{-5} \text{ mM} / fC, P_{Ca} = 0.01, Ca_{min} = 1.0 \cdot 10^{-10} \text{ mM}, \tau_{Ca} = 50.0 \text{ ms}$ |
| $I_{Syn}$ | $g_{Tonic} = 0.31 \text{ nS}, E_{Syn} = -10.0 \text{ mV}, \tau_{Syn} = 5.0 \text{ ms}$ |

For *I*_*Na*_, *I*_*K*_*, I*_*NaP*_, and *I*_*Ca*_, the dynamics of voltage-dependent gating variables *m*_*i*_, and *h*_*i*_ are defined by the following differential equation:

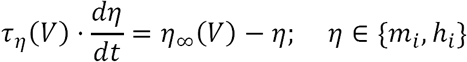

where steady state activation/inactivation *η*_*∞*_ and time constant *τ*_*η*_ are given by:

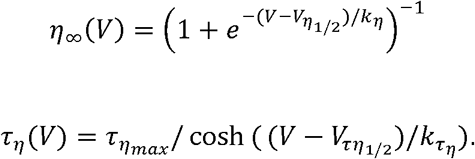

For the voltage-gated potassium channel, steady state activation *m*_*K∞*_*(V)* and time constant *τ*_*mK*_*(V)* are given by:

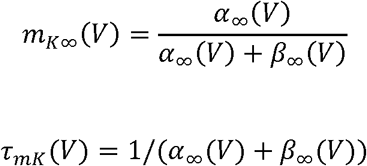

where

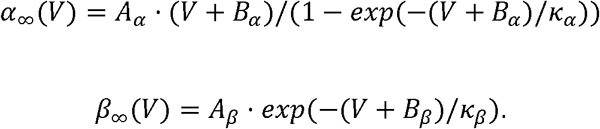

The parameters *V*_*η*_*1/2*, *V*_*τη*_*1/2*, *κ* _*η*_, *κ* _*τη*_*τ*_*η*_*max*, *A*_*α*_, *A*_*β*_, *B* _*α*_, *B*_*β*_, *κ* _*α*_, and *κ* _*β*_ are given in Table 1. *I*_*CAN*_ activation is dependent on the intracellular calcium concentration [*Ca*]_*in*_ and is given by:

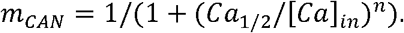

The parameters *Ca*_*l/2*_ and *n*, given in Table 1, represent the half-activation calcium concentration and the Hill Coefficient, respectively.

Calcium enters the neurons through voltage-gated calcium channels (*Ca*_*V*_) and/or synaptic channels (*Ca*_*Syn*_), where a percentage (*P*_*Ca*_) of the synaptic current (*I*_*Syn*_) is assumed to consist of Ca^2+^ ions. A calcium pump removes excess calcium with a time constant *τ*_*Ca*_ and sets the minimum calcium concentration *Ca*_*min*_. The dynamics of [*Ca*]_*in*_ is given by the following differential equation:

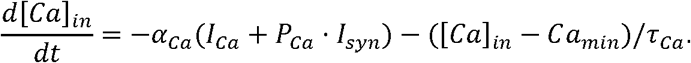

The parameters *α*_*Ca*_ is a conversion factor relating current and rate of change in [*Ca*]_*in*_, see Table 1 for parameter values.

The synaptic conductance of the *i*^th^ neuron 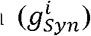 in the population is described by the following equation:

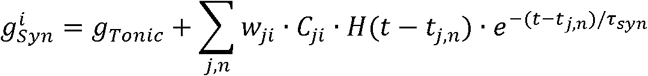

where *w*_*ji*_ is the weight of the synaptic connection from cell *j* to cell *i*, *C* is a connectivity matrix (*C*_*ji*_*= 1* if neuron *j* makes a synapse on neuron *i*, and *C*_*ji*_*= 0* otherwise), *H(.)* is the Heaviside step function, *t* is time, *τ*_*Syn*_ is the exponential decay constant and *t*_*j,n*_ is the time at which an action potential *n* is generated in neuron *j* and reaches neuron *i*.

To account for heterogeneity of neuron properties within the network, the persistent sodium current conductance, 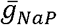, for each neuron was assigned randomly based on a uniform distribution over the range [*0.0,5.0*]*ns* which is consistent with experimental measurements (Rybak et al., 2003; Koizumi & Smith, 2008; Koizumi et al., 2008). The weight of each synaptic connection was uniformly distributed over the range *w*_*ji*_*∈* [*0, W*_*max*_] where *W*_*max*_ ranged from *0.0* to *1.0 ns* depending on the network connectivity and specific simulation. The elements of the network connectivity matrix, *C*_*ji*_, are randomly assigned values of *0* or *1* such that the probability of any connection between neuron *j* and neuron *i* being *1* is equal to the network connection probability *P*_*Syn*_. We varied the connection probability over the range *P*_*Syn*_*∈* [*0.05,1.0*], however, a value of *P*_*Syn*_*= 0.05* was used in most simulations.

### Data Analysis and Definitions

The time of an action potential was defined as when the membrane potential of a neuron crosses *-35mV* in a positive direction. The network activity amplitude and frequency were determined by identifying peaks and calculating the inverse of the interpeak interval in histograms of network spiking. Network histograms of the population activity were calculated as the number of action potentials generated by all neurons per 50 *ms* bin per neuron with units of *spikes/s*. The number of recruited neurons is defined as the peak number of neurons that spiked at least once per bin during a network burst. The average spike frequency of recruited neurons is defined as the number of action potentials per bin per recruited neuron with units of *spikes/s*. The average network resting membrane potential was defined as the average minimum value of *V*_*m*_ in a *500 ms* window following a network burst. The average inactivation of the persistent sodium current at the start of each burst was defined by the maximum of the average value of *h*_*NaP*_ in a 500 *ms* window before the peak of each network burst. The average inactivation of the persistent sodium current at the end of each burst was defined by the maximum of the average value of *h*_*NaP*_ in a 500 *ms* window after the peak of each network burst. Synaptic strength is defined as the number of neurons in the network multiplied by the connection probability multiplied by the average weight of synaptic connections 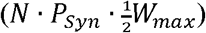. Pacemaker neurons were defined as neurons that continue bursting intrinsically after complete synaptic blockade. Follower neurons were defined as neurons that become silent after complete synaptic blockade. The inspiratory drive potential is defined as the envelope of depolarization that occurs in neurons during the inspiratory phase of the network oscillations (Morgado-Valle et al. 2008).

### Characterization *I*_*CAN*_ in regulating network activity amplitude and frequency in *Ca*_*V*_and *Ca*_*syn*_ Models

To characterize the role of *I*_*CAN*_ in regulation of network activity amplitude and frequency we slowly increased the conductance 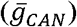 in our simulations from zero until the network transitioned from a rhythmic bursting to a tonic (non-bursting) firing regime. To ensure that the effect(s) are robust, these simulations were repeated over a wide range of synaptic weights, synaptic connection probabilities, and strengths of the intracellular calcium transients from *Ca*_*V*_ or *Ca*_*Syn*_ sources. Changes in network activity amplitude were further examined by plotting the number of recruited neurons and the average action potential frequency of recruited neurons versus 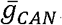.

### Simulated Pharmacological Manipulations

In simulations that are compared with experimental data, both *Ca*_*V*_ and *Ca*_*Syn*_ calcium sources are included. Pharmacological blockade of *I*_*CAN*_ was simulated by varying the conductance, 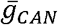 according to a decaying exponential function

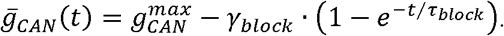

The percent block *γ*_*block*_, decay constant τ_*block*_,and the maximum *I*_*CAN*_ conductance 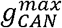 were adjusted to match the experimental changes in network amplitude. The synaptic weight of the network was chosen such that at 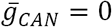 the network activity amplitude was close to 20% of maximum. To reduce the computational time, the duration of *I*_*CAN*_ block simulations was one tenth of the total of experimental durations. For comparison, the plots of normalized change in amplitude and frequency of the simulations were stretched over the same time-period as experimental data. Increasing the simulation time had no effect on our results (data not shown).

### Comparison with Calcium Imaging Data

To allow comparisons with network and cellular calcium imaging data, we analyzed rhythmic calcium transients from our simulations. Single cell calcium signals are represented by [*Ca*]_*i*_. The network calcium signal was calculated as the average intracellular calcium concentration in the network 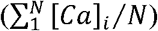.

### Integration Methods

All simulations were performed locally on an 8-core Linux-based operating system or on the high-performance computing cluster Biowulf at the National Institutes of Health. Simulation software was custom written in C++. Numerical integration was performed using the exponential Euler method with a fixed step-size (Δ*t*) of *0.025ms*. In all simulations, the first 50 s of simulation time was discarded to allow for the decay of any initial condition-dependent transients.

## 5. Acknowledgments

This work was supported in part by the Jayne Koskinas Ted Giovanis Foundation for Health and Policy, the Intramural Research Program of the National Institutes of Health (NIH), National Institute of Neurological Disorders and Stroke, and NIH Grants R01 AT008632 and U01 EB021960.

